# *In-silico* docking platform with serine protease inhibitor (SERPIN) structures identifies host cysteine protease targets with significance for SARS-CoV-2

**DOI:** 10.1101/2022.11.18.517133

**Authors:** Joaquín J Rodriguez Galvan, Maren de Vries, Shiraz Belblidia, Ashley Fisher, Rachel A Prescott, Keaton M Crosse, Walter F. Mangel, Ralf Duerr, Meike Dittmann

## Abstract

Serine Protease Inhibitors (SERPINs) regulate protease activity in various physiological processes such as inflammation, cancer metastasis, angiogenesis, and neurodegenerative diseases. However, their potential in combating viral infections, where proteases are also crucial, remains underexplored. This is due to our limited understanding of SERPIN expression during viral-induced inflammation and of the SERPINs’ full spectrum of target proteases. Here, we demonstrate widespread expression of human SERPINs in response to respiratory virus infections, both *in vitro* and *in vivo*, alongside classical antiviral effectors. Through comprehensive *in-silico* docking with full-length SERPIN and protease 3D structures, we confirm known inhibitors of specific proteases; more importantly, the results predict novel SERPIN-protease interactions. Experimentally, we validate the direct inhibition of key proteases essential for viral life cycles, including the SERPIN PAI-1’s capability to inhibit select cysteine proteases such as cathepsin L, and the serine protease TMPRSS2. Consequently, PAI-1 suppresses spike maturation and multi-cycle SARS-CoV-2 replication. Our findings challenge conventional notions of SERPIN selectivity, underscore the power of *in-silico* docking for SERPIN target discovery, and offer potential therapeutic interventions targeting host proteolytic pathways to combat viruses with urgent unmet therapeutic needs.

**SIGNIFICANCE:** Serine protease inhibitors (SERPINs) play crucial roles in various physiological processes, including viral infections. However, our comprehension of the full array of proteases targeted by the SERPIN family has traditionally been limited, hindering a comprehensive understanding of their regulatory potential. We developed an *in-silico* docking platform to identify new SERPIN target proteases expressed in the respiratory tract, a critical viral entry portal. The platform confirmed known and predicted new targets for every SERPIN examined, shedding light on previously unrecognized patterns in SERPIN selectivity. Notably, both key proteases for SARS-CoV-2 maturation were among the newly predicted targets, which we validated experimentally. This underscores the platform’s potential in uncovering targets with significance in viral infections, paving the way to define the full potential of the SERPIN family in infectious disease and beyond.

## INTRODUCTION

Respiratory viruses are group of clinically significant, highly heterogeneous viruses. Despite their differences in particle structure and replication strategies, many rely on a critical group of enzymes: proteases^1^. The origin of these proteases may be host or viral, and the roles they play in viral replication are multifarious^1,2^. The successful application of small molecule protease inhibitors in treating HIV, HCV, and SARS-CoV-2 infections showcased the power of antiviral regimens that target proteases of viral origin^3–5^.

Instead of or in addition to viral-encoded proteases, a number of viruses depend on host proteases to complete their life cycles^6–11^; prominently, for cleavage of fusogenic viral glycoproteins, a process known as viral maturation^6,12,13^. Influenza A viruses (IAV), parainfluenzaviruses, and SARS-CoV-2 all rely on host proteases for maturation of hemagglutinin, fusion, or spike glycoproteins, respectively.^6–12^ Cleavage occurs at specific sites (cleavage sites) to expose the fusion peptide and provide a hinge-like function to enable membrane merging during viral entry. Immature (uncleaved) particles are unable to cause infection. The specificity of host proteases to execute maturation depends on the amino acid motif at the glycoprotein cleavage site ^14^ and the glycoprotein structure adjacent to the cleavage site.^15,16^

The activity of host proteases is controlled at the protein level by host-encoded inhibitors. With over 1,500 members encoded in the genomes of animals, plants, viruses, bacteria, and archaea, SERPINs constitute the largest known family of protease inhibitors.^17^ The mechanism by which SERPINs inhibit target proteases is understood in molecular detail; it results in the formation of irreversible, covalent SERPIN-protease complexes (aka “mousetrap” mechanism, **Supplemental Figure 1**). A SERPIN’s protease specificity is driven mainly by the fit of the SERPIN reactive center loop into the protease catalytic site (**Supplemental Figure 1a**). The discovery of canonical SERPIN target proteases has historically focused on proteases in the blood or the brain^17,18^ and the list of SERPIN targets is likely incomplete. The addition of new SERPIN targets has often occurred incrementally, with discoveries typically arising one at a time during investigations into specific diseases, rather than through systematic discovery approaches. Hence, there is an opportunity to map comprehensive SERPIN target repertoires, particularly within organ-specific proteolytic environments, which remain poorly explored and understood.

**Supplemental Figure 1.**
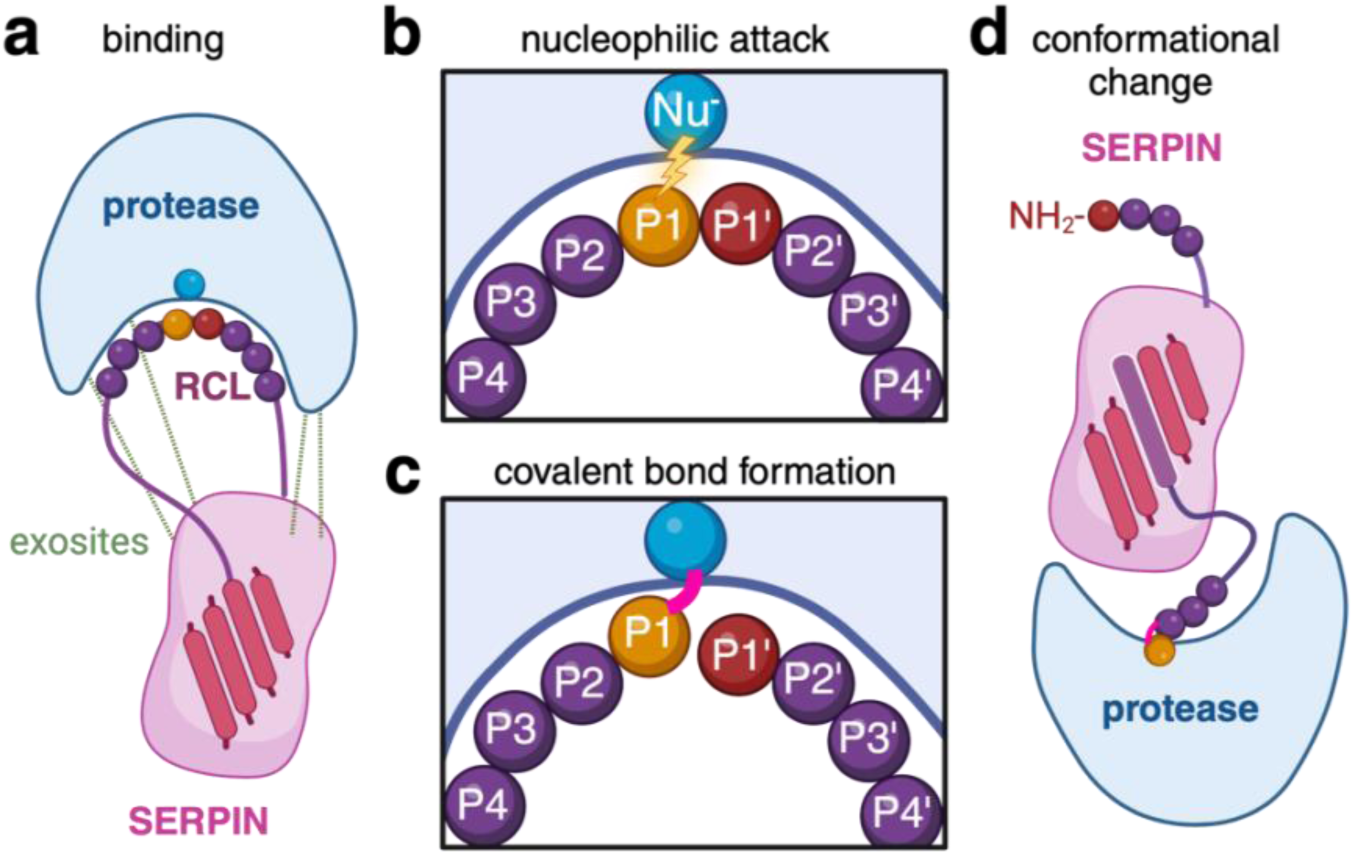
Molecular mode of action for inhibitory SERPINs. **a.** A SERPIN binds to a target protease by inserting its reactive center loop (RCL, purple line) into the protease’s catalytic center. This *binding step* is facilitated by the 3D fit of a given RCL into the catalytic center of the protease and can be enhanced by the formation of secondary binding sites (“exosites”, green dashed lines). **b.** The RCL core sequence (named P4-P4’) mimics the core sequence of the canonical protease substrate. *Nucleophilic attack* by the protease cleaves the SERPIN at the P1-P1’ bond. **c.** Protease and SERPIN form a *covalent complex* (acyl-enzyme intermediate). **d.** The ensuing rapid and significant *conformational change*, where the RCL attached to the protease inserts itself into a ß-sheet center, prompts the formation of a stable inhibitory complex between SERPIN and protease, akin to a “mousetrap”.

To date, three SERPINs have been studied in the context of innate antiviral defense: PAI-1 (encoded by *SERPINE1*) against influenza viruses encoding hemagglutinin H1 and SARS-CoV-2, by impeding the proteolytic maturation of H1 or spike, respectively^19,20^; alpha-1-antitrypsin (encoded by *SERPINA1*) and antithrombin (encoded by *SERPINC1*) against SARS-CoV-2, likely through the inhibition of TMPRSS2, by reducing maturation of spike, although direct inhibition of TMPRSS2 by either SERPIN was not shown^20^. Beyond these investigations, our understanding of the involvement of other SERPIN family members in viral infections remains limited.

Here, we identify SERPINs that are upregulated in airway epithelial cells in response to respiratory viral infections. We devise an *in-silico* docking to predict binding between 10 virus infection-regulated SERPINs and 48 airway proteases based on their 3D structures. All tested SERPINs, despite variations in their specific target repertoires, exhibit predicted binding capabilities with previously unknown protease targets. Notably, these non-canonical targets include crucial proteases involved in respiratory virus infections, including TMPRSS2, a serine protease, and, unexpectedly, several members of the cathepsin (CTS) protease family. Through binding and protease inhibition assays, we confirm that the SERPIN PAI-1 serves as a bona fide direct inhibitor of both TMPRSS2 and cathepsin L (CTSL), with antiviral relevance for SARS-CoV-2 multi-cycle growth. Our findings underscore the importance of expanding the understanding of SERPIN-protease interactions, prove the feasibility of *in-silico* docking strategies for SERPIN target discovery, and highlight the potential of non-canonical interactions as targets for the development of effective antiviral interventions in the context of respiratory viruses.

## RESULTS

### SERPINs are differentially expressed individuals with COVID-19 and in response to respiratory virus infection in a model of the human airway epithelium

While SERPIN expression has been characterized mainly under steady-state conditions, certain SERPINs are upregulated in response to specific stimuli, including interferons, tumor necrosis factors, or tumor growth factors^21–23^, all of which can be produced during viral infections^24–26^. To investigate SERPIN expression in viral infections, we analyzed a published single-cell RNA sequencing (scRNA-seq) dataset of bronchoalveolar lavage fluids (BALF) from individuals with mild or severe COVID-19, as well as uninfected control individuals^27^ (**Figure 1a**). Compared to immune cells such as macrophages, T cells, B cells, and NK cells, we observed a strikingly higher number of upregulated SERPINs in epithelial cells, particularly in samples from individuals with severe COVID-19 (**Figure 1b**). This suggests that airway epithelial cells, which are the major cell type for SARS-CoV-2 replication^28–32^, are the primary source of SERPIN production during SARS-CoV-2 infection.

**Figure 1.**
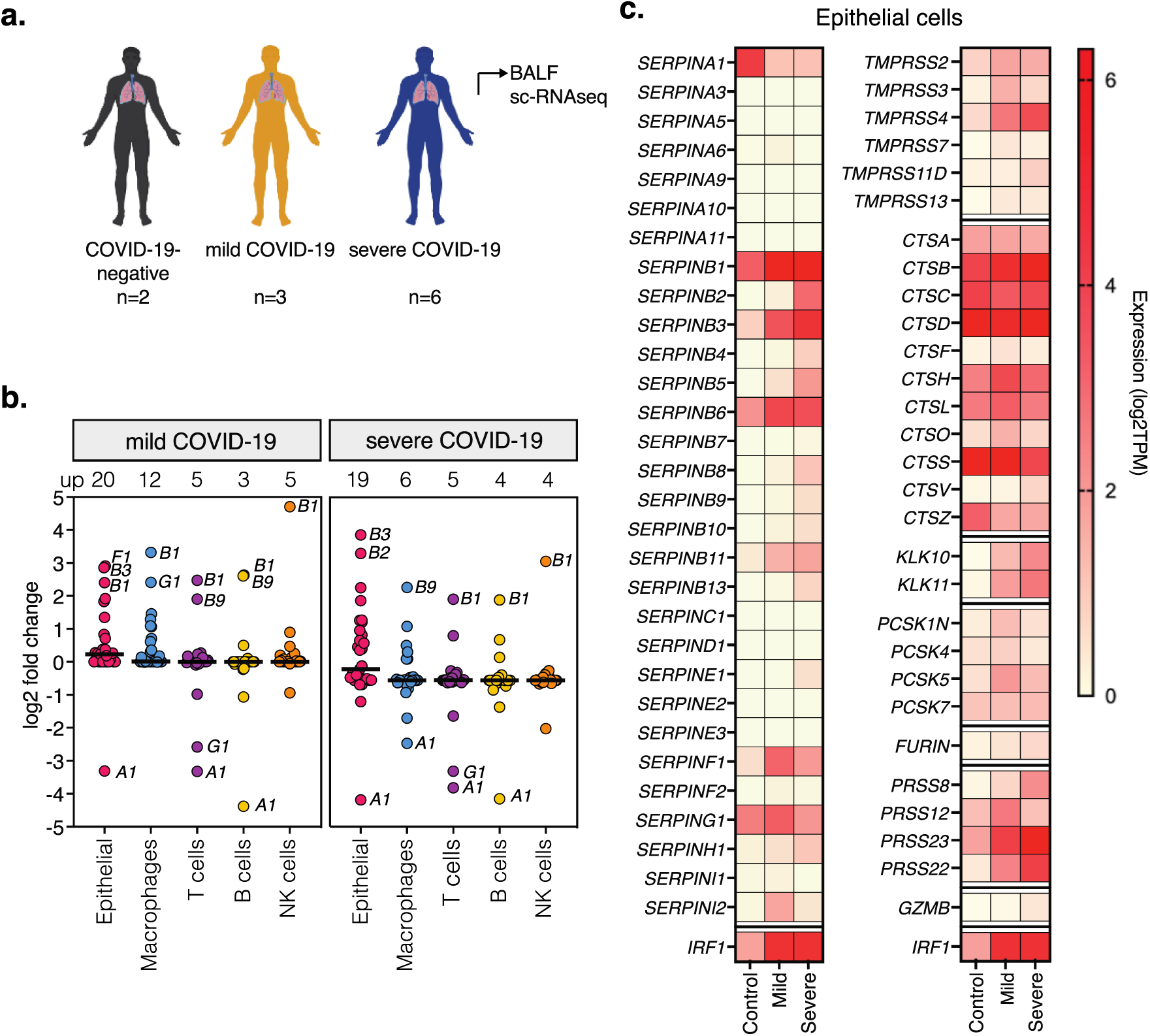
Single-cell transcriptional analysis in bronchioalveolar lavage fluids (BALF) from COVID-19-negative and -positive individuals. **a.** sc-RNAseq data (GSE145926) from BALF of COVID-19-negative (n=2), and COVID-19-positive individuals with mild (n=3) or severe (n=6) symptoms. **b.** Fold-change of SERPIN expression across cell types relative to control individuals. Number of upregulated (>2 fold) SERPINs shown above, with names of most up- or downregulated SERPINs in the graph. Black lines represent average total fold change per group. **c.** Expression of SERPINs, proteases, and prototype interferon-stimulated gene IRF1 in epithelial cells. TPM, Transcripts per million. Only proteases over 0.5 log2TPM in any condition are shown. TMPRSS, transmembrane protease, serine; CTS, cathepsin; KLK, kallikrein; PCSK, pro-protein convertase subtilisin/kexin; PRSS, serine protease; GZMB, granzyme; IRF1, interferon regulatory factor 1.

In epithelial cells, a group of SERPINs (including *SERPINB1*, *SERPINB3*, *SERPINB6*, *SERPING1*) were present at baseline and further upregulated in individuals with COVID-19 (**Figure 1c)**. Others were present at low levels in control individuals and upregulated in COVID-19 individuals (*SERPINB2*, *SERPINB5*, *SERPINB11*, and *SERPINF1).* Yet others were present at low levels irrespective of infection status, and one was downregulated (*SERPINA1*). These observed disparities in expression patterns suggests that specific SERPINs are controlled differentially. Correlation analyses with SERPINs and gene sets involved in inflammatory pathways revealed that mRNA levels for most SERPINs positively correlated with canonical TNF-alpha and interferon-stimulated genes (**Supplemental Figure 2**), indicating that they are upregulated in concert with virus-induced inflammation. Analysis of protease expression in epithelial cells revealed that select proteases were present irrespective of the infection status, while others were upregulated upon infection (**Figure 1c**). This highlights the complexity of the proteolytic landscape produced by airway epithelial cells during SARS-CoV-2 infection.

Given the importance of airway epithelial cells in producing SERPINs during SARS-CoV-2 infection (**Figure 1**), we next used polarized human airway epithelial cultures (HAEC) to study the expression patterns of SERPINs in a controlled environment *in vitro*. The HAEC model closely mimics the human bronchial epithelium, encompassing various cell types (basal, ciliated, and secretory), architectural features, and a secreted extracellular environment that includes mucus and proteases (**Figure 2a, b**)^33^. We infected HAEC with a panel of clinically relevant respiratory viruses (**Figure 2b**), all of which feature proteolytic steps in their life cycles. IFN-beta served as a positive control for expression of interferon-stimulated genes, as evidenced by the IFN-inducible expression of *IFITM3*^34^ (**Figure 2c**).

**Figure 2:**
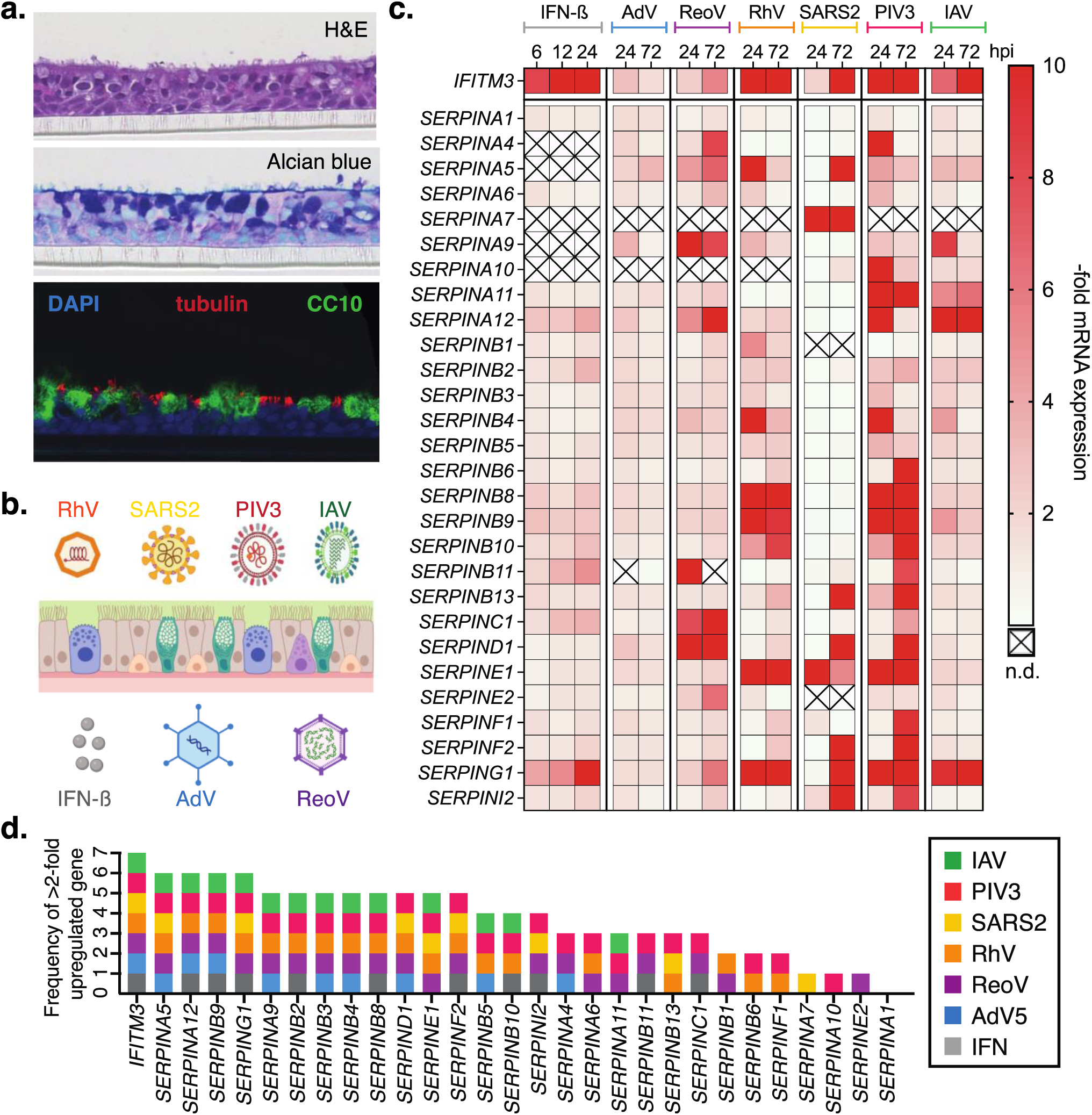
SERPIN expression in virus-infected or interferon-treated human airway epithelial cultures (HAEC). **a.** Analysis of HAEC morphology, with Hematoxylin & Eosin (H&E) staining, Periodic Acid-Schiff (PAS)-Alcian blue staining, and immunofluorescence staining for cell type markers (tubulin in red for ciliated cells, CC10 in green for secretory cells). **b.** Schematic representation of respiratory viruses and a trans-well with polarized HAEC. Apical infection with human rhinovirus A (RhV), influenza A/California/09 H1N1 virus (IAV), human parainfluenzavirus 3 (HPIV3), or SARS-CoV-2 WA-1 (SARS2); basolateral infection/treatment with human adenovirus 5 (AdV5), human reovirus (ReoV), or interferon-beta (IFN-ß). **c.** Total RNA was isolated from lysed cultures at specific time points post-infection, and transcripts were quantified using RT-qPCR. SERPIN and prototype interferon-stimulated gene *IFITM3* mRNA levels shown as fold change over uninfected cultures over time. Data from n=3 biologically independent experiments. **d.** Frequency of >2-fold-upregulated genes from (c) per experimental condition. n.d., not detectable.

**Supplemental Figure 2.**
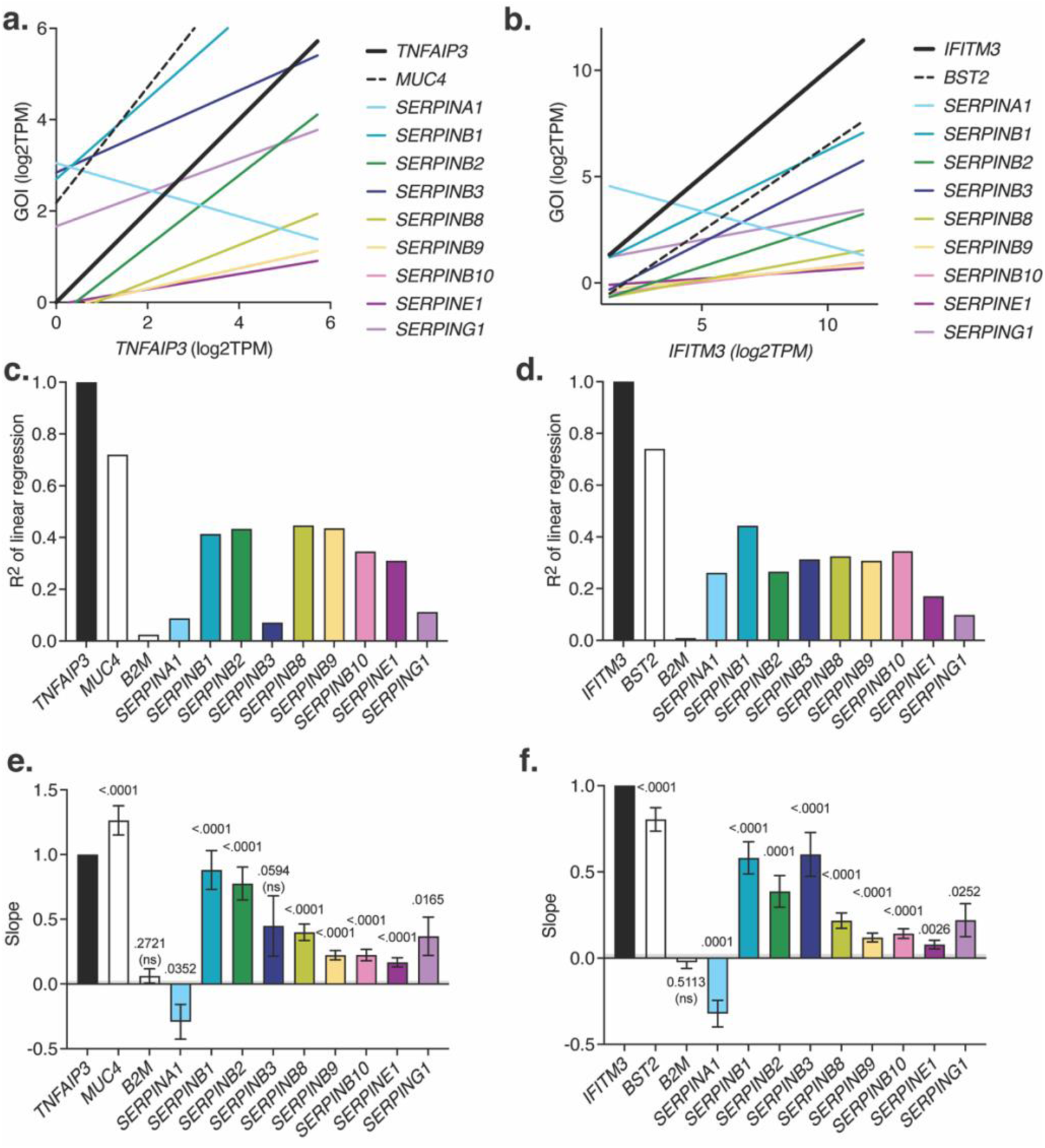
Correlation analysis of SERPIN mRNA with TNF-alpha or type I interferon-induced genes. **a.** Linear regression plot with correlation analysis of select mRNA levels in reference to canonical TNF-alpha-regulated gene TNFAIP3. MUC4, TNF-alpha-regulated gene positive control. **b.** Linear regression plot with correlation analysis of select mRNA levels in reference to canonical type I-interferon-regulated gene IFITM3. BST-2, Interferon-regulated gene positive control. **c.,d.** R2 values of the linear regression plots from a., b., respectively. **e.** Magnitude of slopes of linear regression from (a) and linear regression statistics testing that the slope is significantly non-zero. **f.** Magnitude of slopes of linear regression from (b) and linear regression statistics testing that the slope is significantly non-zero. GOI, gene of interest; ns: not significant

All SERPINs, with the exception of *SERPINA1*, exhibited an upregulation of at least 2-fold under at least one experimental condition (**Figure 2c, d**), suggesting that more SERPINs than previously reported are relevant in the defense to viral infections. A subset of SERPINs was consistently upregulated in response to all six viruses, exemplified by *SERPING1*, suggesting their potential role in a broad antiviral response (**Figure 2c, d**). Other SERPINs exhibited differential expression patterns in response to specific viruses, as seen most notably with *SERPINA7*, *SERPINA10* and *SERPINE2*, indicating distinct regulatory mechanisms and potential selectivity against particular viruses. Different from all other SEPRINs, *SERPINA1* was downregulated upon infection as compared to baseline, suggesting that viral-mediated evasion mechanisms may be at play. We had also found *SERPINA1* expressed at lower levels in BALF of COVID-19 individuals (**Figure 1**). This suggests that *SERPINA1* mRNA expression is downregulated upon SARS-CoV-2 infection, rather than lacking at baseline to allow for SARS-CoV-2 infection, as proposed by others^20^. Finally, only approximately half of the SERPINs upregulated during viral infection were also upregulated upon IFN stimulation (**Figure 2d**), indicating the involvement of other and/or additional gene regulatory mechanisms.

Together, our data demonstrate that most SERPINs are upregulated in response to respiratory virus infections in concert with antiviral effectors, particularly in epithelial cells.

### *In-silico* screen predicts non-canonical SERPIN-protease pairs relevant to viral life cycles

Mapping the full range of proteases targeted by SERPINs remains a challenge, hindering our understanding of SERPIN involvement in human disease. Experimental methods, like SERPIN tagging for pulldown assays and mass spectrometry, require genetic manipulation of model systems. However, many *in-vitro* models fail to replicate the diverse proteolytic landscape observed *in vivo*, limiting the effectiveness of these techniques for target discovery. *In-silico* approaches using motif searches are restricted by known protease motifs and do not integrate structural factors influencing SERPIN-protease interactions.

To overcome this limitation, we developed a computational method to predict 3D interactions between SERPINs and proteases, simulating the binding process depicted in **Supplemental Figure 1a**. Specifically, we employed High Ambiguity Driven protein-protein Docking (HADDOCK), a tool that predicts complex structures, integrating experimental and computational data^35,36^. This method relies on our structural understanding of SERPIN and protease binding directionality, which we use to direct the fit of a SERPIN Reactive Center Loop (RCL) into a protease’s active site. HADDOCK generates multiple iterative conformations of the complex and predicts binding energies for each, with lower scores indicating stronger interactions.

We first evaluated the ability of our approach to distinguish known SERPIN binders from SERPIN non-binders using the well characterized SERPINB1 (**Supplemental Figure 3**). Both the numeric HADDOCK score and visual assessment of docked complexes correctly predicted binding of SERPINB1 to CTSG and Elastase, but not to Granzyme A. As binding is the first step of the SERPIN “mousetrap” mechanism (**Supplemental Figure 1b-d**), we use it as a proxy to screen for predicted SERPIN-protease inhibition.

**Supplemental Figure 3.**
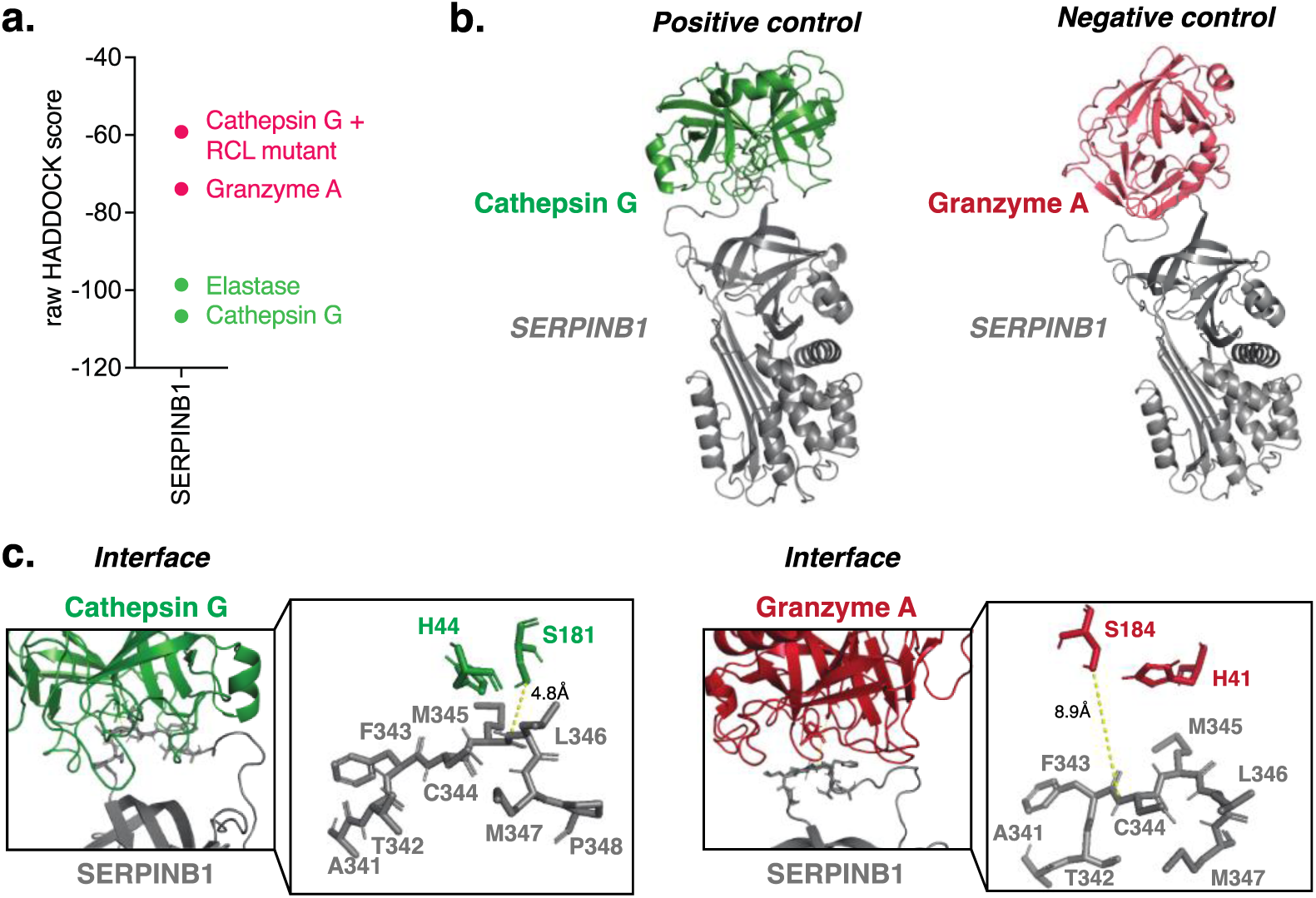
Establishing the *in-silico* screen for identifying SERPIN-protease pairs. **a.** HADDOCK scores reveal predicted binding energies for SERPINB1 with positive controls (Cathepsin G, Elastase) or negative controls (RCL mutant + Elastase, Granzyme A). More negative scores indicate favorable binding. **b.** 3D structure overview featuring SERPINB1 (grey) docked to Cathepsin G (green) or SERPINB1 (grey) docked to Granzyme A (red). **c.** Detailed interface view showing SERPINB1 RCL and protease active sites (Cathepsin G, green, left and Granzyme A, red, right). Zoom-ins reveal residues involved in catalysis. For Cathepsin G, nucleophile S181 and its general base H44 are positioned opposite M345↓L346 with S181 and M345 in close proximity (4.8A). For Granzyme A, nucleophile S184 and its general base H41 are positioned opposite F343↓C344, with S184 and M344 further apart (8.9A).

We next created a panel of 10 SERPINs that we found expressed upon viral infection (**Figure 1, 2)** and 48 potential respiratory target proteases based on the LungMAP human expression database^37^. If available, we gathered the solved 3D structures of each individual protease and SERPIN. If not available, we homology-modeled the 3D structure using Swiss modeler 2.0^38–40^. We mapped protease active sites and SERPIN RCLs by examining individual entries on UniProt (**Figure 3a**). We then docked each of the 10 SERPINs with each of the 48 respiratory proteases, generating a heatmap of normalized HADDOCK scores for each SERPIN-protease pair (**Figure 3b**). Color-coding indicates the energies of the predicted interaction: dark red indicates low HADDOCK scores (favorable binding), and dark grey high HADDOCK scores (unfavorable binding).

**Figure 3:**
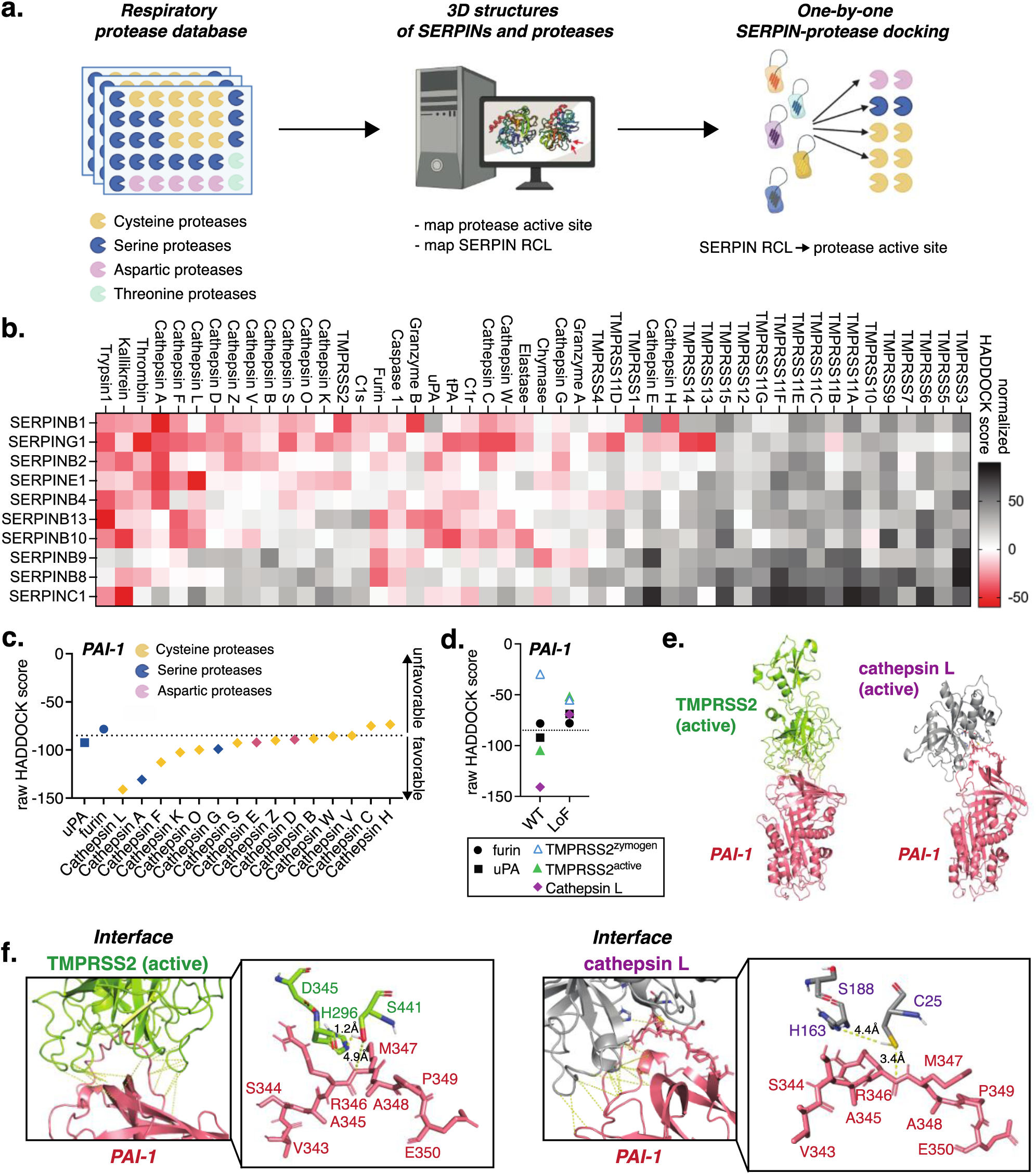
*In-silico* screen of 10 SERPINs against 48 host respiratory proteases. **a.** Schematic of *in-silico*-docking screen workflow. **b.** Heatmap of docking results, with z-scores centered to the mean of control SERPIN-protease pairs and normalized for each SERPIN. The darker the red, the more favorable the binding energies. **c.** Raw HADDOCK scores of PAI-1 with uPA (canonical target), furin (known non-target), and cathepsins. Dotted line separates likely binders from non-binders. **d.** Raw HADDOCK scores of PAI-1 wild type (WT) and loss-of-function (RCL core alanine substitutions, LoF) with uPA, furin, TMPRSS2 zymogen, active TMPRSS2, and active Cathepsin L. **e.** Docking structures of PAI-1 with TMPRSS2 (active) and Cathepsin L. **f.** Details of PAI-1:protease interfaces.

We identified unprecedented patterns in SERPIN selectivity for host proteases involved in respiratory virus infections. A number of SERPINs (i.e., SERPINC1, SERPINB8, and SERPINB9) showed favorable scores with only a few proteases, thereby predicting narrow target specificity, which implies an antiviral profile for viruses relying on those specific proteases. In contrast, other SERPINs exhibited favorable scores for a number of proteases, indicating a broad predicted target range, and positioning them as candidates for blocking redundant proteolytic pathways in viral maturation.

PAI-1 is arguably one of the most-studied SERPINs, with over 15,000 publications^41^. It is also the first-discovered antiviral SERPIN^42^. Our screen revealed favorable scores for PAI-1’s canonical targets uPA/tPA^43^, along with additional previously shown targets trypsin^42^, kallikrein^44^, and thrombin^45^, and, previously suggested but not experimentally shown, TMPRSS2^19,20^ (**Figure 3b**). PAI-1 also docked favorably with a number of CTS, some of which are cysteine proteases (**Figure 3c**). This predicted cysteine protease family binding was unexpected, as PAI-1 was previously thought to have specificity for serine proteases^46^.

Together with TMPRSS2, Cathepsin L (CTSL) is the key protease for SARS-CoV-2 spike S2’ site maturation^47,48^, and we predict both to be targeted by PAI-1, which was previously unknown. We thus further characterized these interactions *in silico*. Mutating the PAI-1 RCL core sequence *in silico* to alanines reduced predicted binding to uPA, TMPRSS2 and CTSL, highlighting the specificity of our docking approach (**Figure 3d**). The PAI-1 score for furin was intermediate, between that of binders and non-binders, and did not change between wild type and loss-of-function PAI-1, suggesting that our approach identifies furin as a non-binder, which is consistent with the literature^42^. PAI-1 only docks favorably with the active form of TMPRSS2 and not with its inactive zymogen, which is in concordance with the SERPIN “mousetrap” mechanism relying on protease activity to initiate a nucleophilic attack.

Visual assessment of the interactions between PAI-1 and TMPRSS2 or CTSL, revealed that the PAI-1 RCL was inserted into the respective proteases’ active sites (**Figure 3e, f**). In the TMPRSS2 complex, the nucleophilic S441 was positioned in close proximity for attack on PAI-1 R346 (**Figure 3f**), thus setting the stage for the next step in the SERPIN “mousetrap” mechanism. R346 is the canonical P1 residue for the PAI-1-uPA inhibitory mechanism. The nucleophilic C25 of CTSL was also positioned in close proximity to canonical P1 PAI-1 R346 (**Figure 3f**), suggesting that this may be the residue that is attacked should the “mousetrap” mechanism occur, which can only be determined experimentally.

In our final *in-silico* validation for PAI-1, we examined published protease target motifs to understand the specificity of proteases for particular substrate motifs, determined by the composition of their specificity pockets within the catalytic center. TMPRSS2 exhibited a preference for an R at P1 and a polar residue at P3, aligning with PAI-1’s core RCL motif (**Supplemental Figure 4**). Conversely, CTSL’s motif flexibility was evident, with no clear consensus motif. However, the PAI-1 RCL core displayed similarities to both the Spike S2’ motif and the minimal motif of CTSL, suggesting potential compatibility with CTSL specificity pockets.

**Supplemental Figure 4.**
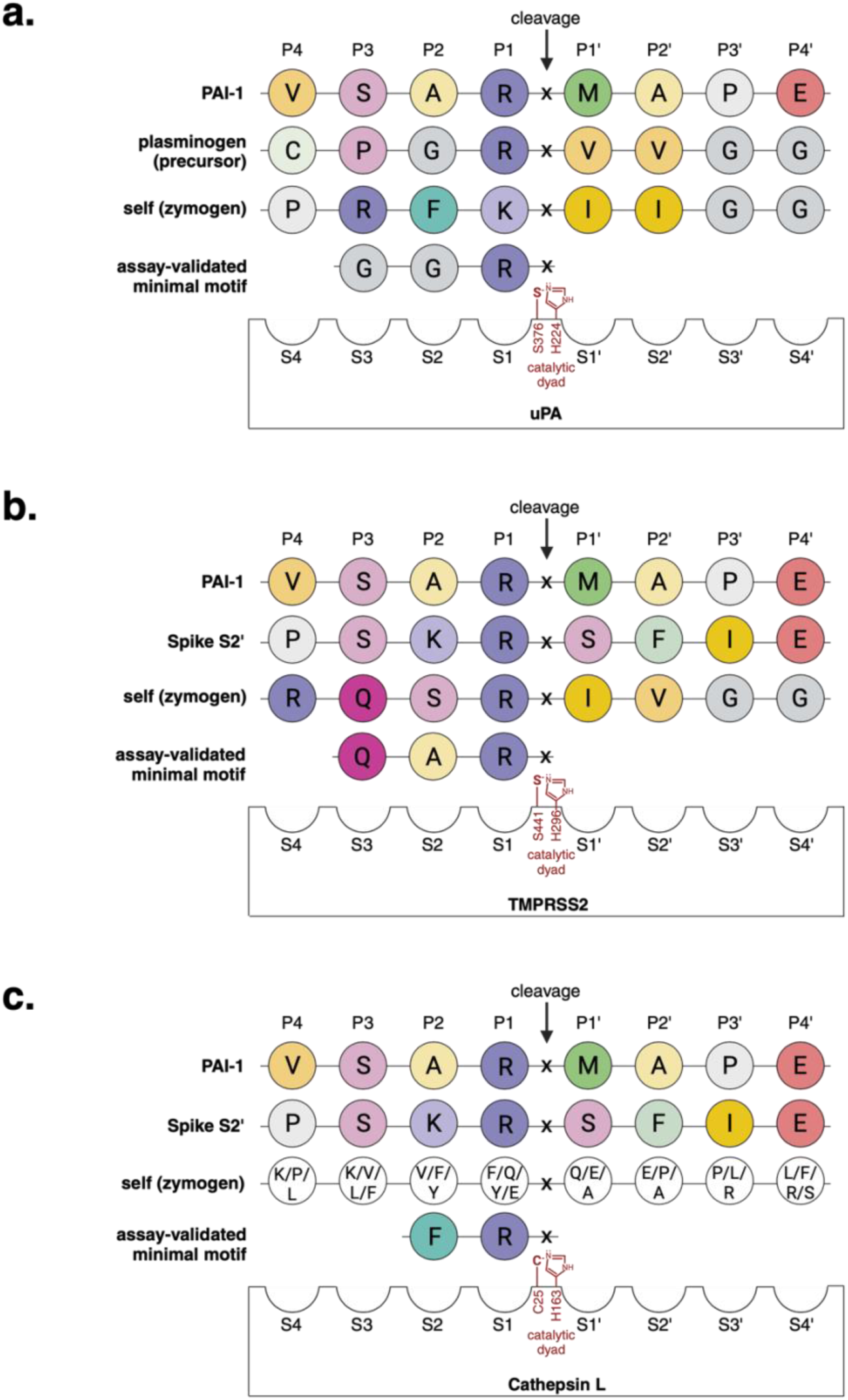
Schematic representation of substrate P4-P4’ motifs opposite proteases’ catalytic pockets S4-S4’ for uPA, TMPRSS2 and Cathepsin L. Amino acids are colored according to their side chain chemistry (Unipro UGENE): basic (R, K) litmus blue with R being more basic and darker; acidic (E, D) litmus red with more acidic being darker; hydrophobic (I, L, V, A), yellow with intensity corresponding to hydrophobic character; sulfur-containing (C, M) green; aromatic (F, Y, W) in teal; polar (N, Q, S, T) magenta/pink with darker coloring for more polarity; non-polar glycine (G) in dark grey and proline (P) in light gray. Assembled from references^49–52^ and UNIPROT.

In summary, our *in-silico* data provide the first comprehensive investigation into novel SERPIN targets within a specific proteolytic environment. We identify numerous potential candidates for antiviral SERPINs warranting further mechanistic investigation. We uncover previously unrecognized patterns in SERPIN selectivity for airway host proteases hijacked by respiratory viruses, such as PAI-1 targeting members of the CTS family, thereby challenging the current understanding of this well-studied SERPIN’s family specificity.

### *In-vitro* assays validate TMPRSS2 and select CTS family members as direct PAI-1 targets with relevance for SARS-CoV-2 maturation

Our *in-silico* screen identified PAI-1 as a predicted inhibitor of both proteases critical for SARS-Cov-2 maturation, TMPRSS2 and CTSL. To determine the inhibitory effect of PAI-1 on TMPRSS2 and CTSL experimentally, we performed fluorometric protease activity assays. Fluorometric protease assays use synthetic reporter-labeled peptide substrates, which undergo changes in absorbance or fluorescence upon cleavage by recombinant proteases. Addition of PAI-1 to the respective protease reactions decreased the activities of all three proteases uPA (a canonical target) as well as TMPRSS2 and CTSL (the predicted targets) in a dose-dependent manner (**Figure 4a-c**).

**Figure 4:**
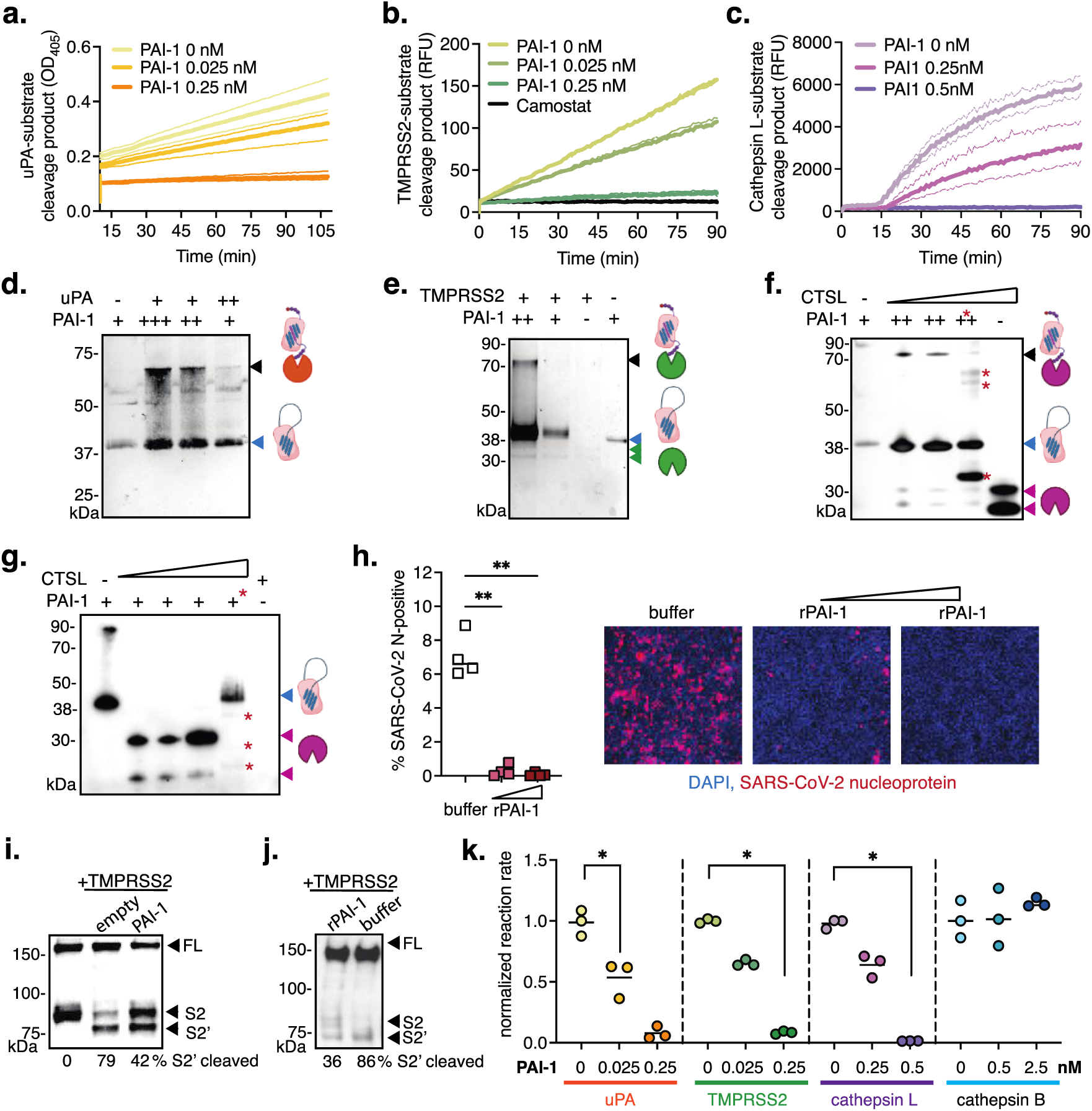
PAI-1 impact on TMPRSS2 and cathepsins and its role in SARS-CoV-2 multi-cycle replication. **a.** uPA protease activity assay with recombinant PAI-1 (Mean ± SD, n=3 replicates). **b.** TMPRSS2 protease activity assay with recombinant PAI-1 or camostat (Mean ± SD, n=3 replicates). **c.** Cathepsin L protease activity assay with recombinant PAI-1 at pH 6.5 (Mean ± SD, n=3 replicates). **d.** SDS-PAGE and silver stain of mixed recombinant active uPA (32 kDa, not visible) and PAI-1 (43 kDa). **e.** SDS-PAGE and silver stain of mixed recombinant active TMPRSS2 (31 kDa) and PAI-1 (43 kDa). **f.,g.** SDS-PAGE and silver stain of mixed recombinant cathepsin L (32 kDa) and PAI-1 (43 kDa) at pH 6.5 (f) or pH 5.5 (g). *Denotes use of PAI-1 inhibitor triplaxinin and PAI-1 cleavage products. **h.** SARS-CoV-2 multi-cycle infection in Calu-3 cells with extracellular addition of recombinant PAI-1 (48 hpi) by high-content microscopy. Statistical analysis by One-way ANOVA and Holm-Sidak’s multiple comparison test, ** p<0.005. Representative images, DAPI (blue, nuclei), SARS-CoV-2 nucleoprotein (red). **i.** BHK cells co-transfected to express SARS-CoV-2 spike, TMPRSS2, and PAI-1. Spike S2 and 2’ band intensities by western blot and densitometry. FL, full-length. **j.** BHK cells co-transfected to express SARS-CoV-2 spike and TMPRSS2, and rPAI-1 or buffer control added to the cell supernatant. Spike S2 and 2’ band intensities by western blot and densitometry. FL, full-length. **k.** Normalized reaction rates for uPA, TMPRSS2, cathepsin L, and cathepsin B in the absence or presence of PAI-1 from *in-vitro* fluorometric assays. Statistical analysis by One-way ANOVA and Kruskal-Wallis test, * p<0.05.

We next tested covalent PAI-1-protease complex formation by incubating recombinant active PAI-1 (rPAI-1) with increasing concentrations of active uPA, TMPRSS2, or CTSL. After incubation, proteins were separated via SDS-PAGE, revealing high molecular weight bands corresponding to predicted complexes (about 75 kDa for PAI-1:uPA, PAI-1:TMPRSS2, and PAI-1:CTSL), indicative of the SERPIN “mousetrap” mechanism (**Figure 4d-f**). Complex formation was specific to conditions with both partners present, with PAI-1:CTSL complexes requiring a pH of 6.5, resembling early endosomal conditions. PAI-1:CTSL high molecular weight complexes did not form in the presence of the PAI-1 inhibitor triplaxinin. Instead, we noted partial digestion of PAI-1 in the presence of triplaxinin, indicating insufficient inhibition of CTSL by PAI-1, consequently rendering PAI-1 susceptible to degradation as a substrate. We also observed partial PAI-1 degradation in the presence of triplaxinin at a pH of 5.5 (**Figure 4 g**). This indicates that, while complexes may not be detectable at levels observable in our experimental setup, inhibition of CTSL by PAI-1 may also occur at this pH.

Next, we determined the antiviral potential of PAI-1 for SARS-CoV-2, given that PAI-1 inhibits both critical maturation proteases: TMPRSS2 plays a crucial role as the spike S2’-cleaving protease for SARS-CoV-2 during viral release (in *cis*) or entry (in *trans*), and CTSL cleaves at SARS-CoV-2 spike at S2’ during endosomal entry, which is the alternative route for SARS-CoV-2 entry. Addition of rPAI to the culture medium of Calu-3 cells significantly reduced SARS-CoV-2 multi-cycle growth compared to carrier control, with near complete inhibition at 6.5 µM (**Figure 4 h**). In fact, we could not detect any spike protein in the supernatants from PAI-1-treated infections, and were thus unable to determine the maturation status of progeny particles.

We next co-transfected BHK cells with spike WA-1 614G, TMPRSS2, and PAI-1 (**Figure 4 i**). While BHK cells express furin for the priming S1/2 cleavage, they lack proteases for the essential S2’ cleavage, detected only with TMPRSS2 overexpression. The presence of PAI-1 reduced S2’ cleavage. Additionally, in a parallel experiment, co-expression of spike WA-1 614G with TMPRSS2 as the S2’-executing protease, along with the addition of recombinant PAI-1 (rPAI-1) to the cell supernatant, led to a reduction in spike S2’ cleavage (**Figure 4 j**). This suggests that PAI-1-mediated TMPRSS2 inhibition contributes to the observed reduction in SARS-CoV-2 multi-cycle replication.

CTSL and CTSB represent the primary cathepsins involved in viral glycoprotein processing^53^. Our docking analysis predicted CTSL as a PAI-1 binder, while CTSB was a predicted as a non-binder (**Figure 3c**), despite both CTS sharing the same minimal consensus target motif, F-R (**Supplemental Figure 4c**). Thus, we assessed CTSB protease reaction rates with and without PAI-1 using fluorometric assays, alongside uPA, TMPRSS2, and CTSL. While PAI-1 reduced the reaction rates of uPA, TMPRSS2, and CTSL, as expected, CTSB remained unaffected (**Figure 4k**), consistent with our *in-silico* predictions. This highlights the significance of the 3D interaction between a SERPIN and a protease in determining specificity, underscoring the power of our *in-silico* docking approach for identifying novel SERPIN targets.

Overall, our findings highlight TMPRSS2 and CTSL as previously unrecognized direct targets of PAI-1. We demonstrate that the cysteine protease family inhibition of CTSL by PAI-1 conforms to the canonical SERPIN “mousetrap” mechanism. Our study broadens our understanding of SERPIN-protease interactions in specific proteolytic contexts, with potential implications beyond infectious disease research.

## DISCUSSION

SERPINs have been extensively investigated in various contexts of human disease, but our understanding of SERPIN expression has mainly been limited to steady-state conditions. Recent sequencing efforts have uncovered the complexity of tissue-specific SERPIN expression^54,55^, and emerging evidence indicates that SERPINs are induced in response to inflammatory cytokines^20,22,23,56,57^. Consistent with these findings, our study demonstrates that SERPIN mRNA expression is upregulated in response to viral infection in both individuals and HAE cultures. Notably, although key SERPINs were found to be upregulated in both, we observed some differences between the two systems. These differences may be attributed to the interplay between respiratory epithelial cells and immune cells, which are lacking in the HAE culture model. For example, it has been demonstrated that macrophage-derived proinflammatory cytokines can stimulate SERPIN expression^58^. Future studies using HAE cultures with exogenously provided cytokines or co-culture systems incorporating immune cells will provide valuable experimental insights on SERPIN transcriptional regulation. Our findings further revealed diverse mRNA expression patterns of specific SERPINs during viral infection, with some SERPINs showing patterns similar to interferon-regulated genes, while others exhibited patterns that were distinct from interferon-stimulated genes and appeared virus-specific. This suggests the involvement of virus-specific regulatory mechanisms beyond interferon signaling, which will be part of future investigations.

Our knowledge of the full array of proteases targeted by the SERPIN family has been limited, hindering a comprehensive understanding of their regulatory potential. Historically, SERPIN target discovery has progressed incrementally, focusing on individual SERPINs or proteases rather than employing systematic discovery approaches. For example, tPA was identified as a PAI-1 target through SDS-PAGE with zymography in plasma samples, searching for SERPIN-protease high molecular weight complexes^59^. More systematic approaches include Co-immunoprecipitation of SERPIN-protease complexes and subsequent partner identification by precipitating either the protease^60^ or the SERPIN^61^. Another strategy is the co-expression of SERPIN variants and select proteases in bacteria, followed by assaying residual protease activity to identify inhibitory variants^62^. Additionally, a recent study generated a RCL mutant library on a known SERPIN background, α1-antitrypsin, tested their inhibitory potential against multiple proteases and, based on the obtained dataset, developed computational algorithms that predict binding denominators within the RCL^63^. This enabled, for the first time, the prediction of how RCLs contribute to target specificity in a comprehensive way. However, information on the 3D structure of both partners, i.e., of the protease’s specificity pocket, and of the formation of exosites between residues outside of the RCL (**Supplemental Figure 1a**) was not considered, and the algorithm relies on data from synthetic SERPINs rather than those occurring in nature. The use of *in-silico* protein-protein docking, which has demonstrated success in identifying compounds that modulate enzymatic reactions, including those involving proteases, could overcome these limitations. Yet, *in-silico* docking is constrained by the need for detailed knowledge on the binding directionality of the two partners, such as guiding the docking of a small molecule inhibitor into the catalytic center of a protease where it is functionally relevant. Fortuitously, the well-understood and conserved nature of SERPIN-protease interactions has enabled us to leverage *in-silico* protein-protein docking for SERPIN target discovery in our study, by guiding the SERPIN RCL into a protease’s catalytic center. Our approach provides a quantitative, numeric score for predicted SERPIN-protease interaction energies and, additionally, offers the structural assessment of bound complexes, thereby ensuring appropriate fit, the visualization of potential exosites, and the approximate prediction of P1 residues. Importantly, it transcends the reliance on protease motif similarities alone, as exemplified by our correct identification of CTSL but not CTSB as a PAI-1 target, despite those two CTS sharing the same minimal substrate motif. This was possible because our approach considers the overall SERPIN RCL fit into the 3D structure of the protease’s specificity pocket. In all, our approach extends beyond the identification of SERPIN specificity patterns and allows for molecular predictions of how SERPINs bind to non-canonical proteases.

One limitation of our screening approach is that it does not consider SERPIN cofactors. For example, SERPINC1 requires heparin as a cofactor to mediate its interaction with thrombin, its canonical target, in a bridge-like manner^54^. To overcome this limitation, the crystal structure of the SERPINC1-heparin-thrombin complex can serve as a template for modeling^64^. Another potential limitation is that our docking methodology does not account for the subcellular localization of SERPINs and their targets, e.g., within specific compartments or the extracellular space. For instance, PAI-1, a secreted SERPIN, exhibits anti-protease activity in the extracellular space^65^. The TMPRSS2 zymogen is membrane-bound initially and inactive, and its active form is released into the extracellular space upon auto-cleavage^66^. Thus, it is feasible that PAI-1 and the TMPRSS2 active form come into contact and interact in the extracellular space. Indeed, our previous research has demonstrated that the exit of PAI-1 from cells is crucial for providing antiviral protection against influenza virus spread^19^. Although CTSL is typically found within endosomes^67^, it can also be secreted^68,69^, and the receptor-mediated uptake of extracellular PAI-1 via endocytosis and the described anti-protease activity of PAI-1 within endosomes have been shown^70,71^. This implies the potential for PAI-1 to interact with CTSL either extracellularly or within endosomes. Our *in-vitro* findings indicate PAI-1-CTSL complex formation preferentially at a pH resembling that of early endosomes. Thus, while our docking method does not account for SERPIN and target protease localization, integrating literature and conducting additional experiments, such as altering the extracellular presence of protease or SERPIN^20,42^, or redirecting SERPINs to different compartments^72^, can overcome this limitation.

Our study marks the first comprehensive exploration of SERPIN targets within a specific proteolytic context, by docking SERPINs with a panel of 48 proteases expressed in the respiratory epithelium. We observed that most members of the TMPRSS family were not identified as targets of the SERPINs tested, except for TMPRSS13 and 14, predicted for SERPING1 only; TMPRSS1, predicted for SERPINB1 only; and TMPRSS11D, TMPRSS4, and TMPRSS2, predicted to interact with multiple SERPINs. Conversely, certain host proteases such as trypsin, plasma kallikrein, and CTSA were predicted to bind with nearly all tested SERPINs, indicating potential sensitivity to inhibition by multiple SERPINs. Our findings highlight numerous potential candidates for antiviral SERPINs, warranting further mechanistic exploration beyond this study. We also unveil previously unnoticed patterns in SERPIN selectivity, exemplified by PAI-1 targeting members of the CTS family, thereby challenging the prevailing understanding of this extensively studied SERPIN’s specificity. CTS play pivotal roles in various viral diseases, regulating viral antigen presentation, facilitating viral release through extracellular matrix remodeling, modulating cell death, and contributing to viral glycoprotein maturation in viruses such as Ebola virus, Nipah virus, Hendra virus, and SARS-CoV-2^53^. Notably, both CTSB and CTSL are the main CTS known to provide viral glycoprotein maturation. The observation that PAI-1 exclusively inhibits CTSL and not CTSB suggests that other SERPINs with predicted specificity for CTSB (SERPINB1, SERPINB2, and SERPING1) may be needed to bridge this gap. The discovery of PAI-1 targeting CTSL, previously unknown, along with our demonstration of PAI-1’s direct inhibition of TMPRSS2 for the first time in this study, implicates the inhibition of both primary (TMPRSS2) and auxiliary (CTSL) proteases in SARS-CoV-2 maturation. Our findings indicate a functional link between extracellular PAI-1 levels and the control of SARS-CoV-2 spread, potentially explaining the underlying mechanism behind other reports linking the presence of the PAI-1 single nucleotide polymorphism rs6092 to increased disease severity in COVID-19^73^. Moreover, the discovery of PAI-1 targeting CTSL may have implications beyond infectious diseases, as this pair has been found to be upregulated concurrently in the metastasis of various cancers^74–76^, suggesting a functional rather than merely correlative aspect of disease outcome.

Viral glycoproteins have evolved to exploit multiple host proteases redundantly for maturation, as seen in SARS-CoV-2 spike’s processing by TMPRSS2 and CTS L. Thus, targeting host proteases like TMPRSS2 with Camostat individually has not been fully effective in antiviral therapy^7,77,78^. Similarly, innate host defenses must block a combination of proteases to effectively inhibit viral glycoprotein maturation. Our screen predicts that most SERPINs inhibit multiple proteases. SERPINB1, a member of the B clade, demonstrated extensive predicted protease binding, encompassing various CTSs, including some previously identified^54,79^, as well as trypsin and furin. Another broad-spectrum candidate is SERPINB13, which showed favorable scores throughout the tested CTS, in accordance with its known role as a CTS inhibitor^80^, but also scored favorably with furin, which was previously unknown. It must be noted that none of the tested SERPINs demonstrated comprehensive inhibitory activity against all known pro-viral host protease classes, suggesting a need for a combination of SERPINs to effectively inhibit viral replication by targeting complementary proteolytic pathways in the human airway epithelium.

The investigation of host proteases as targets in antiviral treatments has been a key focus in drug development^81^. Our foundational study reveals the antiviral potential of host-encoded protease inhibitors and suggests key proteolytic pathways that can be targeted for antiviral therapies by mirroring the action of antiviral SERPINs. Another possibility is the expression or administration of full-length recombinant SERPINs with redirected specificity^82^ or short SERPIN RCL peptides^83^. An alternative option is the design of small molecules stabilizing SERPIN-protease binding^17^. These approaches aim to replicate the SERPIN mechanism while mitigating potential side effects often associated with full-length wild type SERPINs, which retain their impact on unwanted host proteases and can thus lead to adverse effects when overexpressed. Beyond SARS-CoV-2, similar approaches as in this study could be undertaken to discover antiviral SERPIN candidates relevant for proteolytic landscapes in other viral entry portals, such as the gut, and for emerging viruses with predicted protease reliance based on their glycoprotein cleavage sites.

## MATERIAL AND METHODS

### COVID19 Patient sc-RNASeq data acquisition and analysis

Publicly available COVID-19 patient sc-RNASeq data were obtained from the Gene Expression Omnibus database under the accession code GSE145926 ^27^. Aligned and normalized transcript per million (TPM) values were obtained from the integrated ToppCell web server ^84^. Frequencies, and log2 fold changes were calculated from the normalized TPM values by the healthy controls. Of note, monocytes, innate lymphoid cells, neutrophils, and dendritic cells, although detected in COVID-19 individuals, were not detected in uninfected individuals’ BALF, thereby preventing the determination of fold changes for these cell types. Upregulated SERPINs were defined as those expressed to >2-fold higher levels in COVID-19 individuals compared to uninfected controls. Plots were generated in GraphPad Prism v9.

### Generation of polarized human airway epithelial cultures

Differentiation of hTert-immortalized bronchial basal progenitor cells into pseudo-stratified human airway epithelial cultures was performed as stated in the following publication ^85^. Briefly, Bci-NS1.1 cells (obtained from the Laboratories of Drs. Matthew Walters and Ronald Crystal, Cornell University) were seeded into Pneumacult™-Ex Plus Medium (StemCell) and passaged at least two times before plating (7×10^4^ cells/well) on rat-tail collagen type-1 coated permeable transwell membrane supports (6.5mm; Corning Inc) to generate human airway epithelial cultures (HAE). HAE were maintained in Pneumacult™- Ex Plus Medium until confluent, then grown at an air-liquid interface with Pneumacult™-ALI Medium in the basal chamber for approximately 3 weeks to form differentiated, polarized cultures that resemble *in vivo* pseudo-stratified mucociliary epithelium.

### Viral infection of human airway epithelial cultures and transcript determination by RT-qPCR

For infection with influenza A/California/07/2009 virus (BEI resources, NR-13663), HPIV3-GFP (kind gift of Dr. Peter Collins), Rhinovirus A16 (kind gift of Dr. Ann Palmenberg), and SARS-CoV-2 WA-1 (BEI resources) cultures were washed twice with room-temperature PBS, then virus was added apically (5×10^6^ PFU for IAV, 1.12×10^7^ FFU for PIV, 0.39 TCID_50_ for Rhinovirus, and 1.35E5 PFU for SARS-CoV-2) in 50 µl of PBS and incubated for one hour at 37°C for IAV and PIV, and at 4°C for 90 minutes for Rhinovirus. Viruses were aspirated and HAE cultures were washed apically once with room-temperature PBS, then incubated at 37°C until endpoint. For infection with human Reovirus T3 SA (kind gift of Dr. Terrence Dermody) and Adenovirus 5-GFP (kind gift of Dr. Patrick Hearing), transwells were placed inverted in a sterile 5cm petri dish bottom and all remaining basal media was aspirated from the basal side of the transwells. 25 µl of each virus was added to the basal side of the transwells (1.3×10^8^ PFU for Reovirus and 8.85×10^5^ PFU for Adenovirus) and incubated for one hour at 37°C for Adenovirus and room temperature for Reovirus. Viruses were aspirated from the basal side of the transwells, and transwells were placed back into ALI media and incubated at 37°C until endpoint. At each endpoint, transwell membranes were submerged in 375 µl Buffer RLT (Qiagen) and cell lysate was frozen at -80°C for future RNA extraction.

RNA was isolated using the Qiagen RNeasy kit following manufacturer’s instructions. Extracted RNA was then quantified via NanoDrop, and diluted to correct for yield difference. Relative mRNA levels, normalized to housekeeping gene RPS11, were determined by RT-qPCR (SuperScript III First Strand Synthesis System, Life Technologies and PowerUP SYBR Green Master Mix, Thermo Fisher Scientific). Primers used are listed in **Supplemental Table 1**.

### SARS-CoV-2 infection of Calu-3 cells under addition of extracellular rPAI-1

We used 5E4 PFU SARS-CoV-2 WA-1 per culture in a black, clear bottom 96-well plate from corning. Target rPAI-1 levels were chosen to reflect physiological concentrations, at 25 or 12.5 ng/ml, respectively^42^. Virus was diluted in DMEM and supplemented with rPAI-1 to the concentrations mentioned above diluted in PBS, or buffer (50mM NaH_2_PO_4_, 100mM NaCl, 1mM EDTA, pH 6.6). Cells were incubated with virus dilutions rocking for 2h at 37C, then removed. rPAI-1 or buffer dilutions to the same concentration were replenished in after, diluted in DMEM (DMEM, 10%FBS, 1X Penicillin/Streptomycin, 1X Non-essential amino acids) for the course of the infection until fixing. Cells were fixed in 10% Formalin overnight and used for immunofluorescence for detection of SARS-CoV-2 Nuclecapsid protein positive cells.

### Assembly of Protease and SERPIN database and structure modeling

Protein structures were obtained directly from the Protein Data Bank (PDB) or modeled by homology using swiss PDB modeler ^38–40^. If obtained from the PDB, structures were refined by removing solvents, co-factors, small molecules, and co-crystallized atoms. Selection of proteases for later docking was based on the LungMAP database ^84^ of human airway expressed proteases and protease families, mainly prioritizing those expressed to >10 Log2TPM on epithelial cells of donors. For proteases expressed as zymogens, we derived their active forms from the UniProt server ^86^ and modeled the active forms. Active sites and SERPIN RCLs were mapped on to refined structures using the UniProt server^86^.

### SERPIN-protease docking using HADDOCK v2.4

Docking of SERPIN-protease complexes were performed using the HADDOCK v2.4 server ^35,87^. Selection of primary residues to perform docking was based on the residues specified on UniProt as the reactive center loop’s 8 critical residues found to dictate SERPIN specificity - P4-P4’ for SERPINs and the catalytic triad within the protease active site for proteases. Secondary residues, also taken into accont while guiding the docking, were automatically selected within the range of a 6.5 Angstrom radius from each of the primary residues. Furthermore, the full RCL was given full flexibility, starting at the poly-alanine segment and ending at the PF loop end motif. Upon submission and finish of an entry on the HADDOCK v2.4 server, we evaluated the resulting complex position and specifically selected complexes that had RCLs fit inside the protease active site, if no such complex was found, the protease was removed from the screen. We then used the score of the visually fitting complexes and used it to guide our first initial optimization and calculate thresholds for the z-scores. Structures of the complexes were then saved and visualized on PyMoL v2.0 for distance calculation and deeper examination of the positioned residues (Figure 4) ^88^.

### Statistical analysis of raw HADDOCK scores and conversion into normalized HADDOCK scores

To define the thresholds of SERPIN-protease docking, Haddock scores were determined from at least two positive and two negative control proteases for each SERPIN, known to bind or not bind SERPINs *in vitro,* respectively. For each SERPIN, average HADDOCK scores were calculated separately for the positive and negative controls, and the mean of both averaged scores served as the in-silico threshold of SERPIN-protease interaction. In cases where negative controls were not available, we generated *in-silico* mutants in the SERPIN reactive center loop (RCL) by converting the P4-P4’ residues to alanines. The mutated SERPIN structures were generated using Swiss modeler 2.0^38–40^, and then docked to positive control proteases. Swiss modeler was preferred over AlphaFold for *in-silico* mutagenesis, as the AlphaFold algorithm disregards structural changes caused by introduced mutations. The SERPIN-specific HADDOCK thresholds were used to normalize the full data set of HADDOCK scores and to generate Z-scores, i.e., the thresholds were set to zero. Violin plots were generated using GraphPad Prism v.9, and clustered heatmaps of the normalized, Z-scored data were generated using the ComplexHeatmap package in program R v.4.1.2 ^89^.

### Antibodies, recombinant proteins, and inhibitors

To treat cells with rPAI-1 we used recombinant active human PAI-1 (carrying stabilizing mutations N150K, K154T, Q319L and M354I, Abcam) in experiments on human airway epithelial cultures and in *in-vitro* complex formation assays. rTMPRSS2 was obtained from Biomatik (RPC24995), rCTSL from R&D Systems (952-CY-010), rCTSB from R&D Systems (953-CY-010) and recombinant active urokinase plasminogen activator from (Abcam). Triplaxtinin (PAI-039, Axonmedchem) was used as PAI-1 inhibitor for *in-vitro* assays at 10 uM. To visualize spike in Western Blots we used anti SARS-CoV-2 Spike S2 (1A9) mouse monoclonal antibody (ThermoFisher, MA5-35946) at 1:1000 diluted in 3%BSA in TBS-T (0.01% Tween). For visualization of SARS-CoV-2 infected cells via detection of N protein in Calu-3 cells we used anti SARS-CoV-1/2 Nucleocapsid Mouse anti-N clone (Cell Signaling, 1C7C7) 1:1000 50uL/well.

### SERPIN-protease complex formation assays

Reactions were set up in PCR tubes containing a constant amount of 0.5 ug or 0.1 ug active recombinant PAI-1, co-incubated with a protease or not, respectively, per reaction. Reactions were conducted in the following buffers: TMPRSS2 and uPA reactions used 10mM Tris-HCl, 200mM NaCl, 10mM EDTA, pH 7.4; Cathepsin L used 50 mM MES, 5 mM DTT, 1 mM EDTA, pH 6.5 and pH 5.5 for early and late endosome mimics, respectively. Protease concentrations were used as follows: TMPRSS2, 0.1 ug; Cathepsin L, 0.25ug, and 1ug when incubated alone; uPA, 1 ug and 2 ug. Reaction tubes were incubated in a thermocycler for 10 min at 25C and 30 min at 37 C, consecutively. When applicable, Triplaxtinin was used at 10µM.

### Fluorometric protease activity assays

Commercially available recombinant protease activity kits were used to quantify activity and inhibition of the mentioned proteases in the presence of rPAI-1. TMPRSS2 kit from BPS Bioscience (#78083) was used and instructions were followed from the kit for kinetic measurements, rPAI-1 was used at 0nM, 0.025nM and 0.25nM. Cathepsin L kit from BPS Bioscience (#79591) was used and instructions were followed, with the only modification of adjusting the pH of the 1X reaction buffer solution to pH 6.5 to maximize PAI-1 activity. This kit was also used for Cathepsin B screening only by changing the enzyme. Both enzymes were used at a final concentration of 0.5nM and rPAI-1 was added to a final concentration of 0.25nM and 0.5nM, or 0.5nM and 2.5nM for Cathepsin L and B, respectively. Fluorescence or absorbance was measured over time using a SpectraMax M5 model by Molecular Devices (for TMPRSS2) and Cathepsins B and L, and uPA substrate emitted fluorescence or changes in absorbance was measured in kinetic mode using EnVision plate reader (#1041590 EnVision). Reaction slopes were calculated by fitting the raw time-course values curve to a simple linear regression model. Slopes were then normalized to enzyme alone to obtain the relative rate.

## Supporting information

Supplemental files are integrated in the manuscript file

## ACKNOWLEDGEMENTS

We thank all members of the Dittmann lab for critical input on data and manuscript. We also thank Dr. Terrence Dermody for the kind gift of Reovirus T3 SA, Dr. Ann Palmenberg for the kind gift of Rhinovirus A16, Dr. Peter Collins for the kind gift of PIV3-GFP, and Dr. Patrick Hearing for the kind gift of Adenovirus 5-GFP. Research was supported by the following grants from NIH/NIAID: R01AI143639 and R21AI139374 to M.D. as well as T32AI100853. Work was further supported by The Vilcek Institute of Graduate Biomedical Sciences, and by NYU Grossman School of Medicine Startup funds.

---

**Supplemental Table 1.**
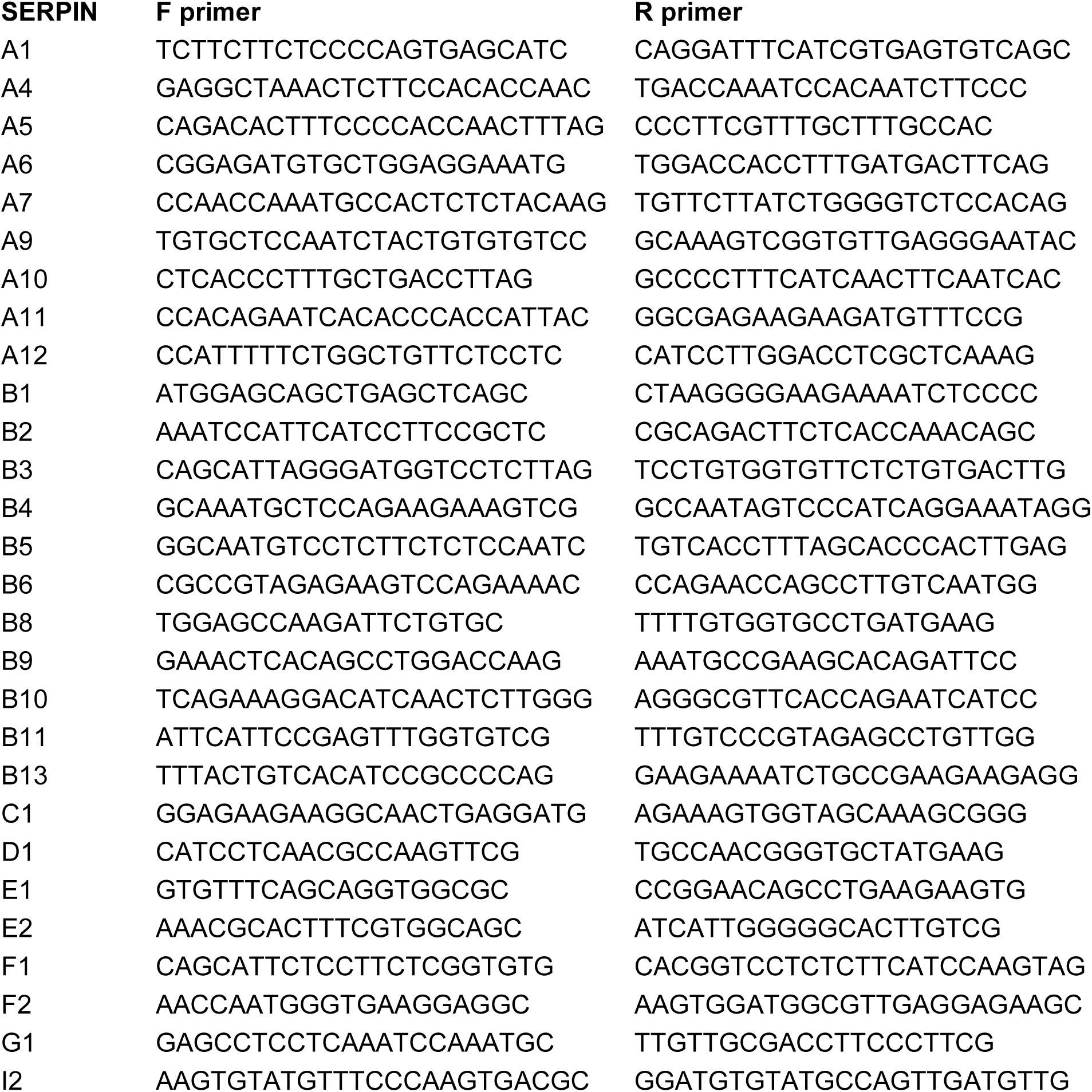

