## Supplemental files are integrated in the manuscript file for "*In-silico* docking platform with serine protease inhibitor (SERPIN) structures identifies host cysteine protease targets with significance for SARS-CoV-2"

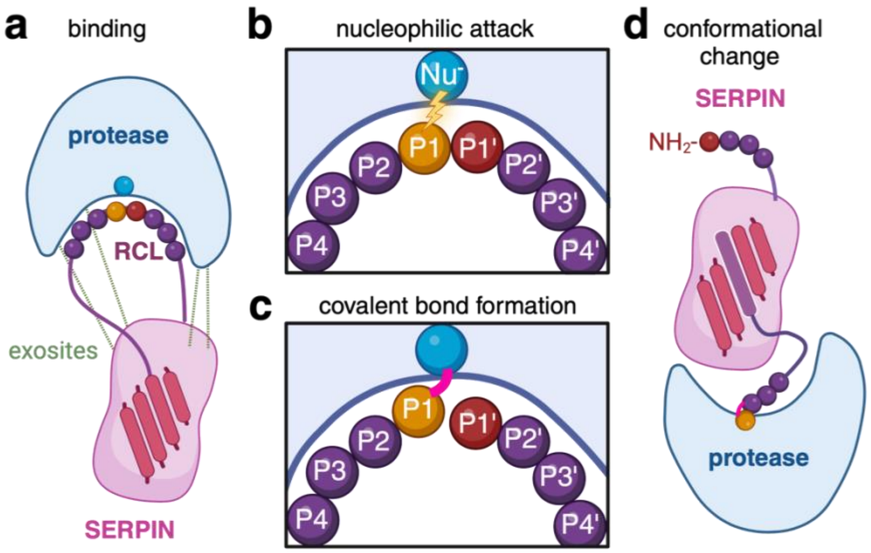


**Supplemental Figure 1. Molecular mode of action for inhibitory SERPINs. a.** A SERPIN binds to a target protease by inserting its reactive center loop (RCL, purple line) into the protease's catalytic center. This *binding step* is facilitated by the 3D fit of a given RCL into the catalytic center of the protease and can be enhanced by the formation of secondary binding sites ("exosites", green dashed lines). **b.** The RCL core sequence (named P4-P4') mimics the core sequence of the canonical protease substrate. *Nucleophilic attack* by the protease cleaves the SERPIN at the P1-P1' bond. **c.** Protease and SERPIN form a *covalent complex* (acyl-enzyme intermediate). **d.** The ensuing rapid and significant *conformational change*, where the RCL attached to the protease inserts itself into a ß-sheet center, prompts the formation of a stable inhibitory complex between SERPIN and protease, akin to a "mousetrap".

### RESULTS

#### SERPINs are differentially expressed individuals with COVID-19 and in response to respiratory virus infection in a model of the human airway epithelium.

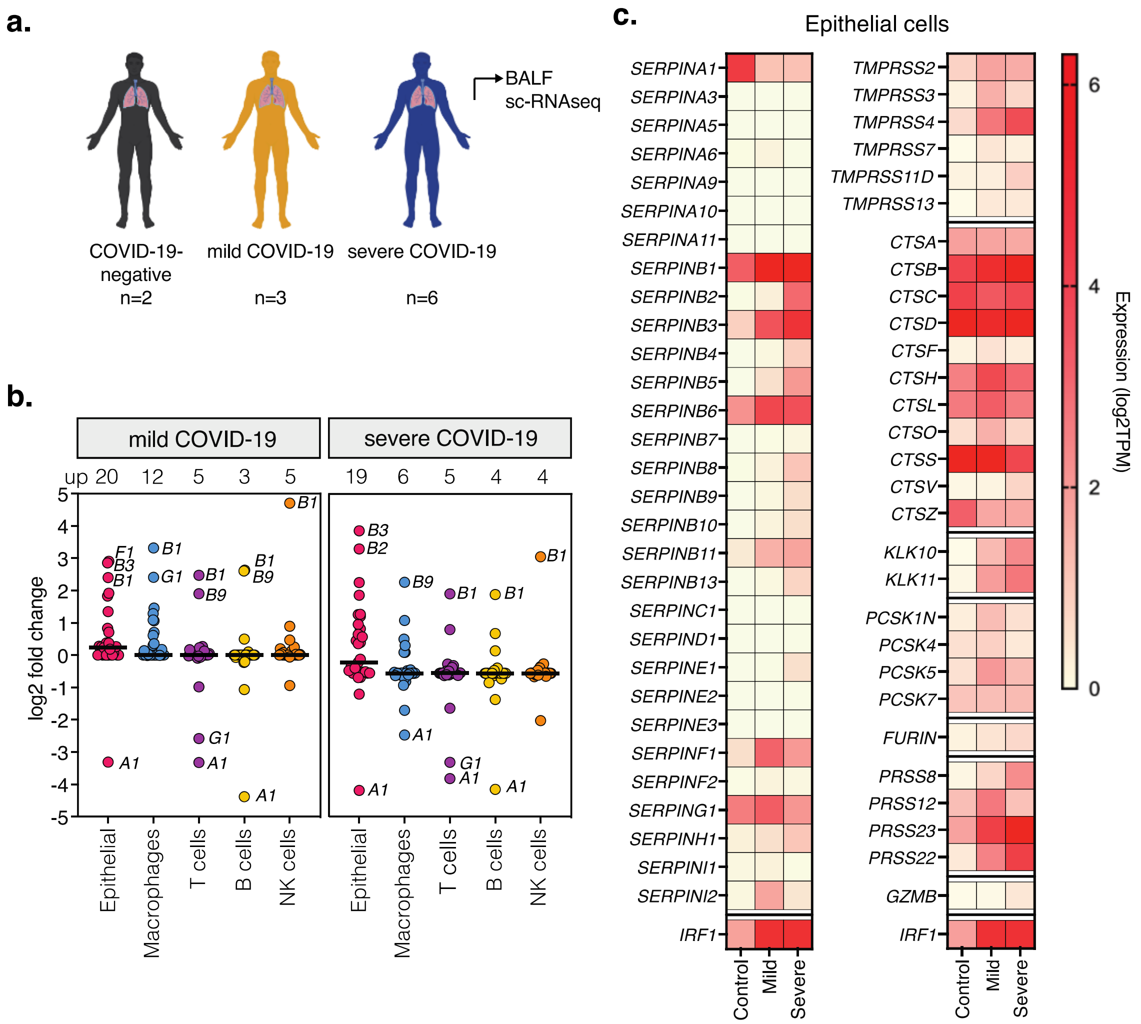


**Figure 1. Single-cell transcriptional analysis in bronchioalveolar lavage fluids (BALF) from COVID-19-negative and -positive individuals. a.** sc-RNAseq data (GSE145926) from BALF of COVID-19-negative (n=2), and COVID-19-positive individuals with mild (n=3) or severe (n=6) symptoms. **b.** Fold-change of SERPIN expression across cell types relative to control individuals. Number of upregulated (>2 fold) SERPINs shown above, with names of most up- or downregulated SERPINs in the graph. Black lines represent average total fold change per group. **c.** Expression of SERPINs, proteases, and prototype interferon-stimulated gene IRF1 in epithelial cells. TPM, Transcripts per million. Only proteases over 0.5 log2TPM in any condition are shown. TMPRSS, transmembrane protease, serine; CTS, cathepsin; KLK, kallikrein; PCSK, pro-protein convertase subtilisin/kexin; PRSS, serine protease; GZMB, granzyme; IRF1, interferon regulatory factor 1.

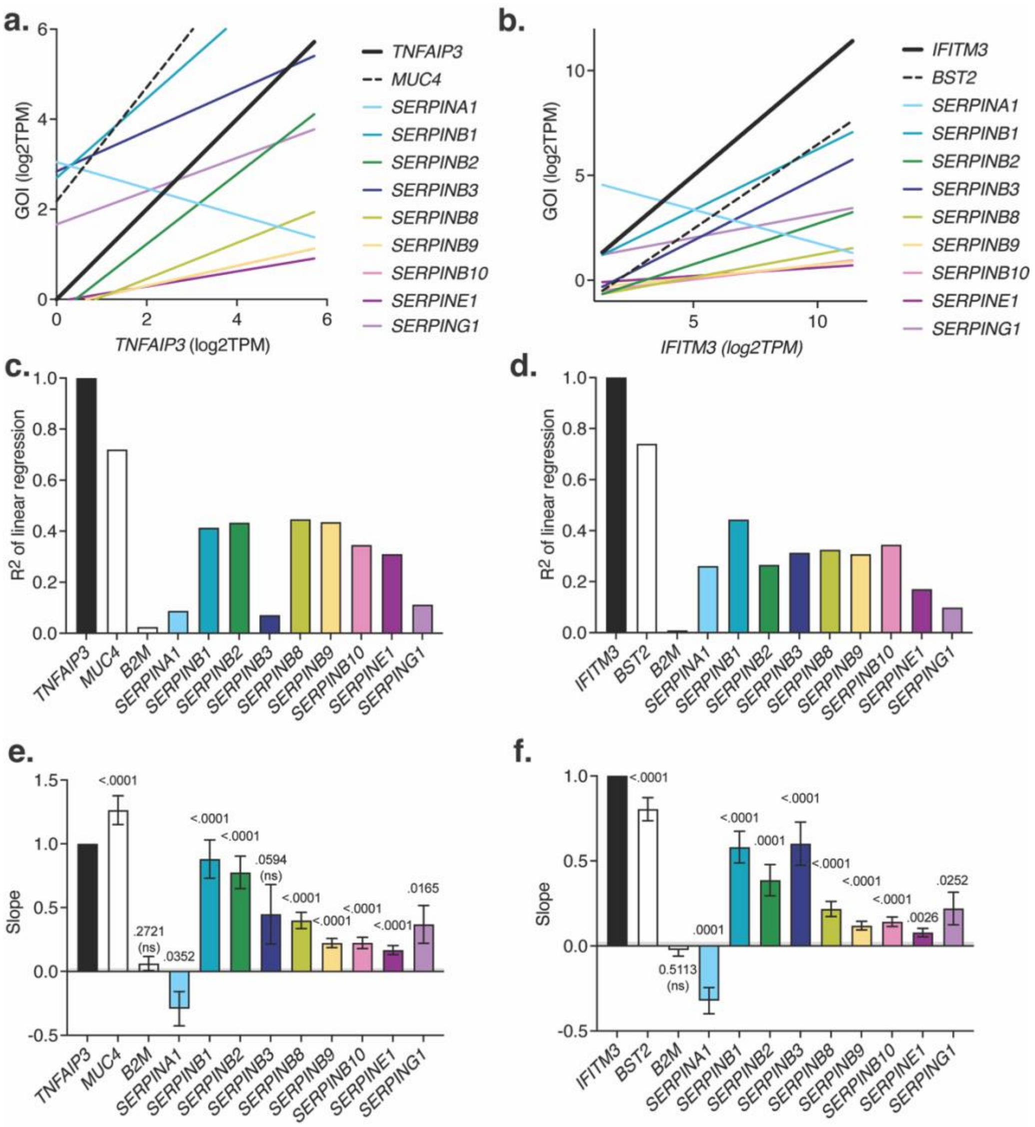


**Supplemental Figure 2**. **Correlation analysis of SERPIN mRNA with TNF-alpha or type I interferon-induced genes. a.** Linear regression plot with correlation analysis of select mRNA levels in reference to canonical TNF-alpha-regulated gene TNFAIP3. MUC4, TNF-alpha-regulated gene positive control. **b.** Linear regression plot with correlation analysis of select mRNA levels in reference to canonical type I-interferon-regulated gene IFITM3. BST-2, Interferon-regulated gene positive control. **c.,d.** R2 values of the linear regression plots from a., b., respectively. **e.** Magnitude of slopes of linear regression from (a) and linear regression statistics testing that the slope is significantly non-zero. **f.** Magnitude of slopes of linear regression from (b) and linear regression statistics testing that the slope is significantly non-zero. GOI, gene of interest; ns: not significant

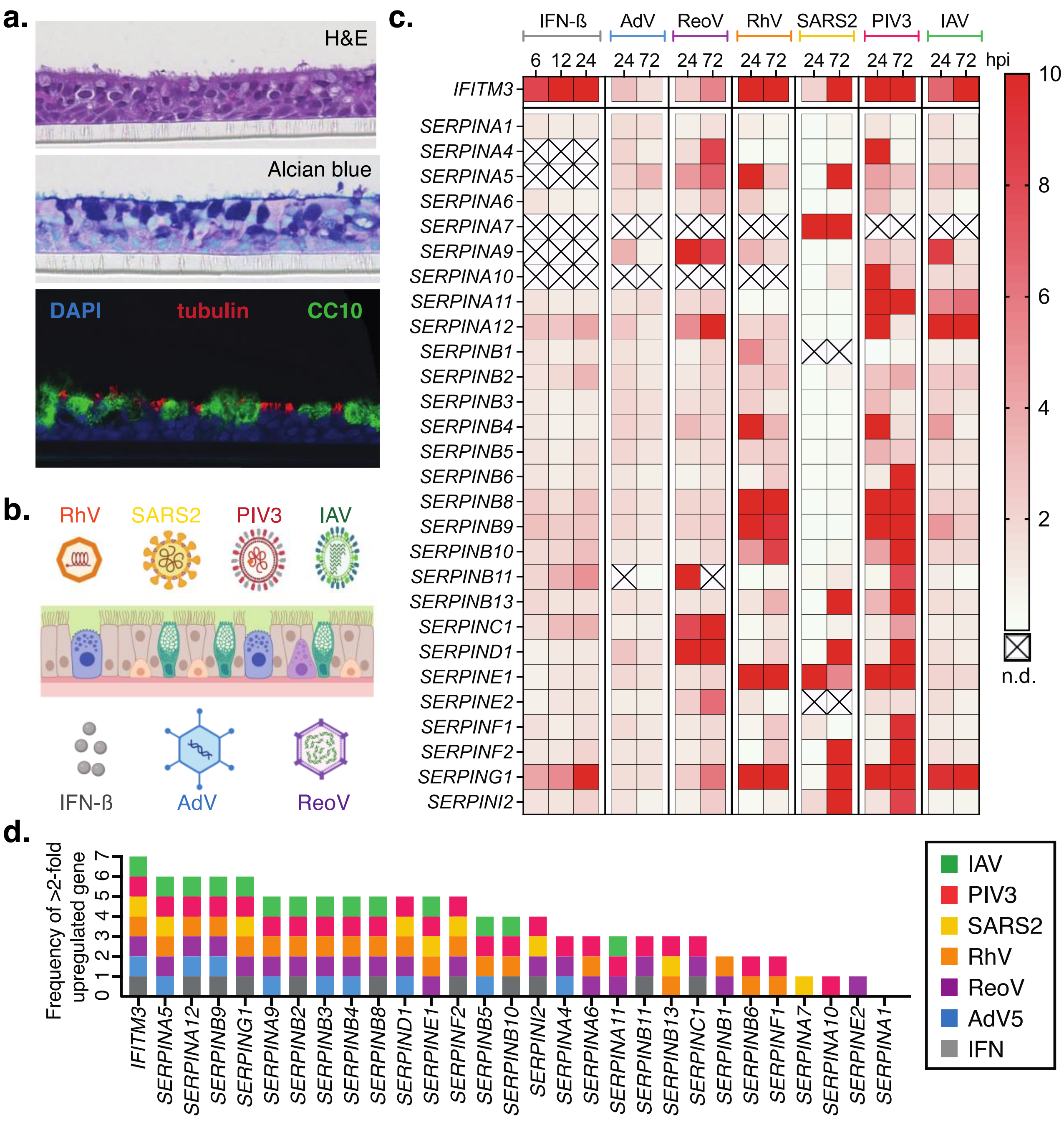


**Figure 2: SERPIN expression in virus-infected or interferon-treated human airway epithelial cultures (HAEC). a.** Analysis of HAEC morphology, with Hematoxylin & Eosin (H&E) staining, Periodic Acid-Schiff (PAS)-Alcian blue staining, and immunofluorescence staining for cell type markers (tubulin in red for ciliated cells, CC10 in green for secretory cells). **b.** Schematic representation of respiratory viruses and a trans-well with polarized HAEC. Apical infection with human rhinovirus A (RhV), influenza A/California/09 H1N1 virus (IAV), human parainfluenzavirus 3 (HPIV3), or SARS-CoV-2 WA-1 (SARS2); basolateral infection/treatment with human adenovirus 5 (AdV5), human reovirus (ReoV), or interferon-beta (IFN-ß). **c.** Total RNA was isolated from lysed cultures at specific time points post-infection, and transcripts were quantified using RT-qPCR. SERPIN and prototype interferon-stimulated gene *IFITM3* mRNA levels shown as fold change over uninfected cultures over time. Data from n=3 biologically independent experiments. **d.** Frequency of >2-fold-upregulated genes from (c) per experimental condition. n.d., not detectable.

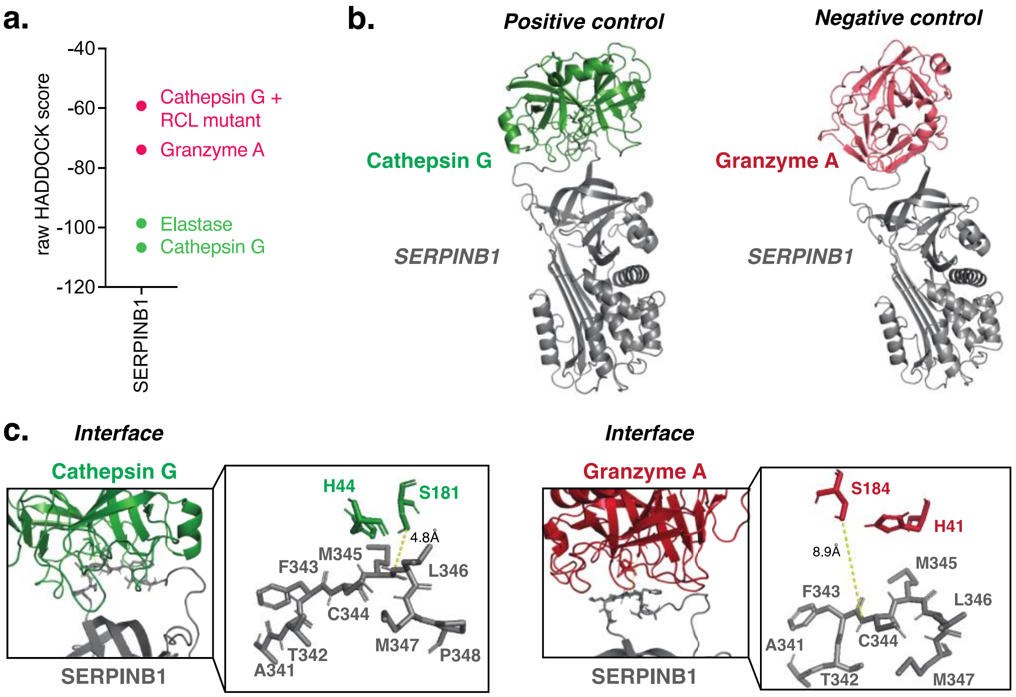


**Supplemental Figure 3. Establishing the *in-silico* screen for identifying SERPIN-protease pairs. a.** HADDOCK scores reveal predicted binding energies for SERPINB1 with positive controls (Cathepsin G, Elastase) or negative controls (RCL mutant + Elastase, Granzyme A). More negative scores indicate favorable binding. **b.** 3D structure overview featuring SERPINB1 (grey) docked to Cathepsin G (green) or SERPINB1 (grey) docked to Granzyme A (red). **c.** Detailed interface view showing SERPINB1 RCL and protease active sites (Cathepsin G, green, left and Granzyme A, red, right). Zoom-ins reveal residues involved in catalysis. For Cathepsin G, nucleophile S181 and its general base H44 are positioned opposite M345↓L346 with S181 and M345 in close proximity (4.8A). For Granzyme A, nucleophile S184 and its general base H41 are positioned opposite F343↓C344, with S184 and M344 further apart (8.9A).

We first evaluated the ability of our approach to distinguish known SERPIN binders from SERPIN non-binders using the well characterized SERPINB1 (**Supplemental Figure 3**). Both the numeric HADDOCK score and visual assessment of docked complexes correctly predicted binding of SERPINB1 to CTSG and Elastase, but not to Granzyme A. As binding is the first step of the SERPIN "mousetrap" mechanism (**Supplemental Figure 1b-d**), we use it as a proxy to screen for predicted SERPIN-protease inhibition.

We next created a panel of 10 SERPINs that we found expressed upon viral infection (**Figure 1, 2)** and 48 potential respiratory target proteases based on the LungMAP human expression database[37]. If available, we gathered the solved 3D structures of each individual protease and SERPIN. If not available, we homology-modeled the 3D structure using Swiss modeler 2.0[38-40]. We mapped protease active sites and SERPIN RCLs by examining individual entries on UniProt (**Figure 3a**). We then docked each of the 10 SERPINs with each of the 48 respiratory proteases, generating a heatmap of normalized HADDOCK scores for each SERPIN-protease pair (**Figure 3b**). Color-coding indicates the energies of the predicted interaction: dark red indicates low HADDOCK scores (favorable binding), and dark grey high HADDOCK scores (unfavorable binding).

We identified unprecedented patterns in SERPIN selectivity for host proteases involved in respiratory virus infections. A number of SERPINs (i.e., SERPINC1, SERPINB8, and SERPINB9) showed favorable scores with only a few proteases, thereby predicting narrow target specificity, which implies an antiviral profile for viruses relying on those specific proteases. In contrast, other SERPINs exhibited favorable scores for a number of proteases, indicating a broad predicted target range, and positioning them as candidates for blocking redundant proteolytic pathways in viral maturation.

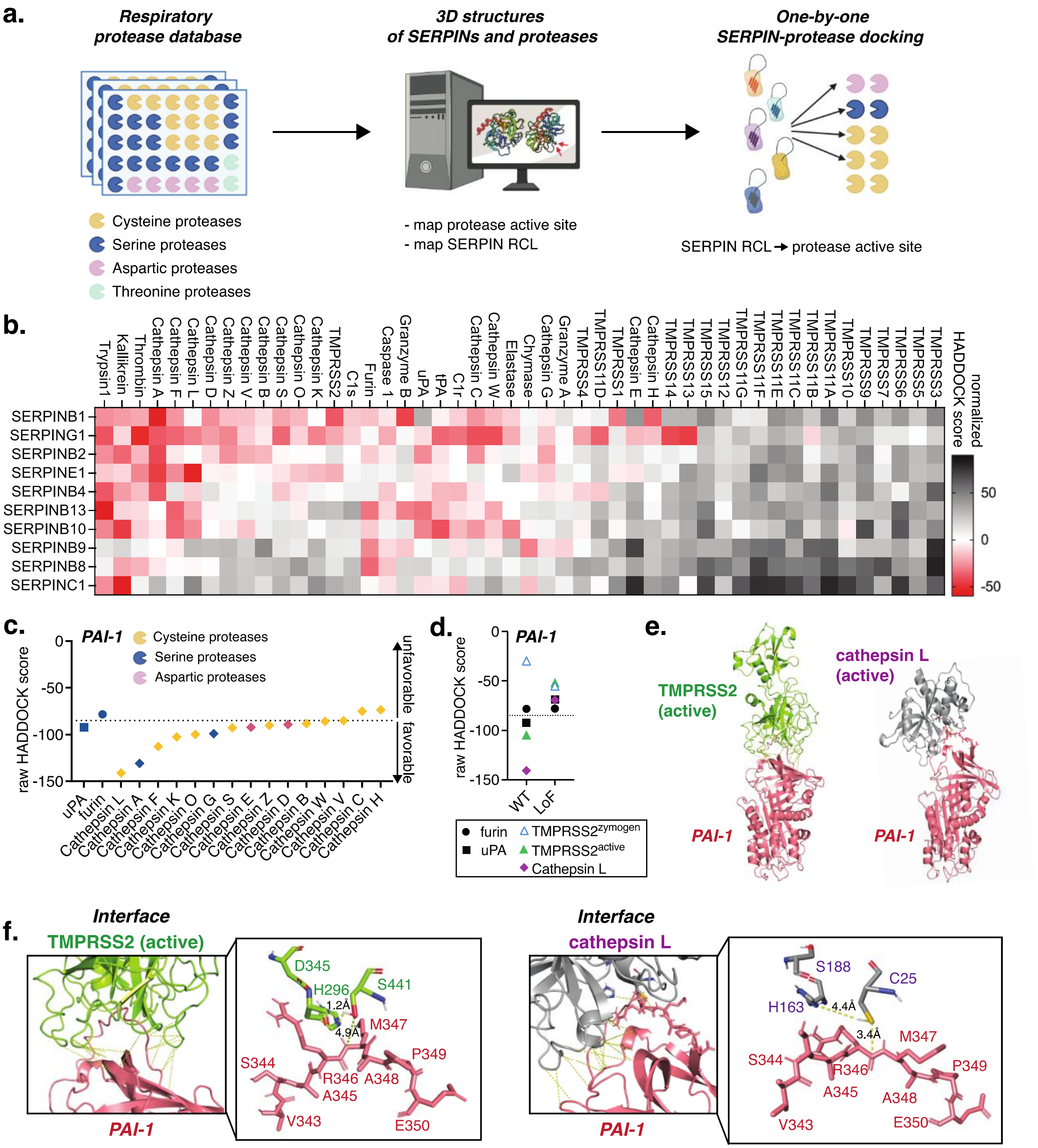


**Figure 3: *In-silico* screen of 10 SERPINs against 48 host respiratory proteases. a.** Schematic of *in-silico*-docking screen workflow. **b.** Heatmap of docking results, with z-scores centered to the mean of control SERPIN-protease pairs and normalized for each SERPIN. The darker the red, the more favorable the binding energies. **c.** Raw HADDOCK scores of PAI-1 with uPA (canonical target), furin (known non-target), and cathepsins. Dotted line separates likely binders from non-binders. **d.** Raw HADDOCK scores of PAI-1 wild type (WT) and loss-of-function (RCL core alanine substitutions, LoF) with uPA, furin, TMPRSS2 zymogen, active TMPRSS2, and active Cathepsin L. **e.** Docking structures of PAI-1 with TMPRSS2 (active) and Cathepsin L. **f.** Details of PAI-1:protease interfaces.

PAI-1 is arguably one of the most-studied SERPINs, with over 15,000 publications[41]. It is also the first-discovered antiviral SERPIN[42]. Our screen revealed favorable scores for PAI-1's canonical targets uPA/tPA[43], along with additional previously shown targets trypsin[42], kallikrein[44], and thrombin[45], and, previously suggested but not experimentally shown, TMPRSS2[19,20] (**Figure 3b**). PAI-1 also docked favorably with a number of CTS, some of which are cysteine proteases (**Figure 3c**). This predicted cysteine protease family binding was unexpected, as PAI-1 was previously thought to have specificity for serine proteases[46].

Together with TMPRSS2, Cathepsin L (CTSL) is the key protease for SARS-CoV-2 spike S2' site maturation[47,48], and we predict both to be targeted by PAI-1, which was previously unknown. We thus further characterized these interactions *in silico*. Mutating the PAI-1 RCL core sequence *in silico* to alanines reduced predicted binding to uPA, TMPRSS2 and CTSL, highlighting the specificity of our docking approach (**Figure 3d**). The PAI-1 score for furin was intermediate, between that of binders and non-binders, and did not change between wild type and loss-of-function PAI-1, suggesting that our approach identifies furin as a non-binder, which is consistent with the literature[42]. PAI-1 only docks favorably with the active form of TMPRSS2 and not with its inactive zymogen, which is in concordance with the SERPIN "mousetrap" mechanism relying on protease activity to initiate a nucleophilic attack.

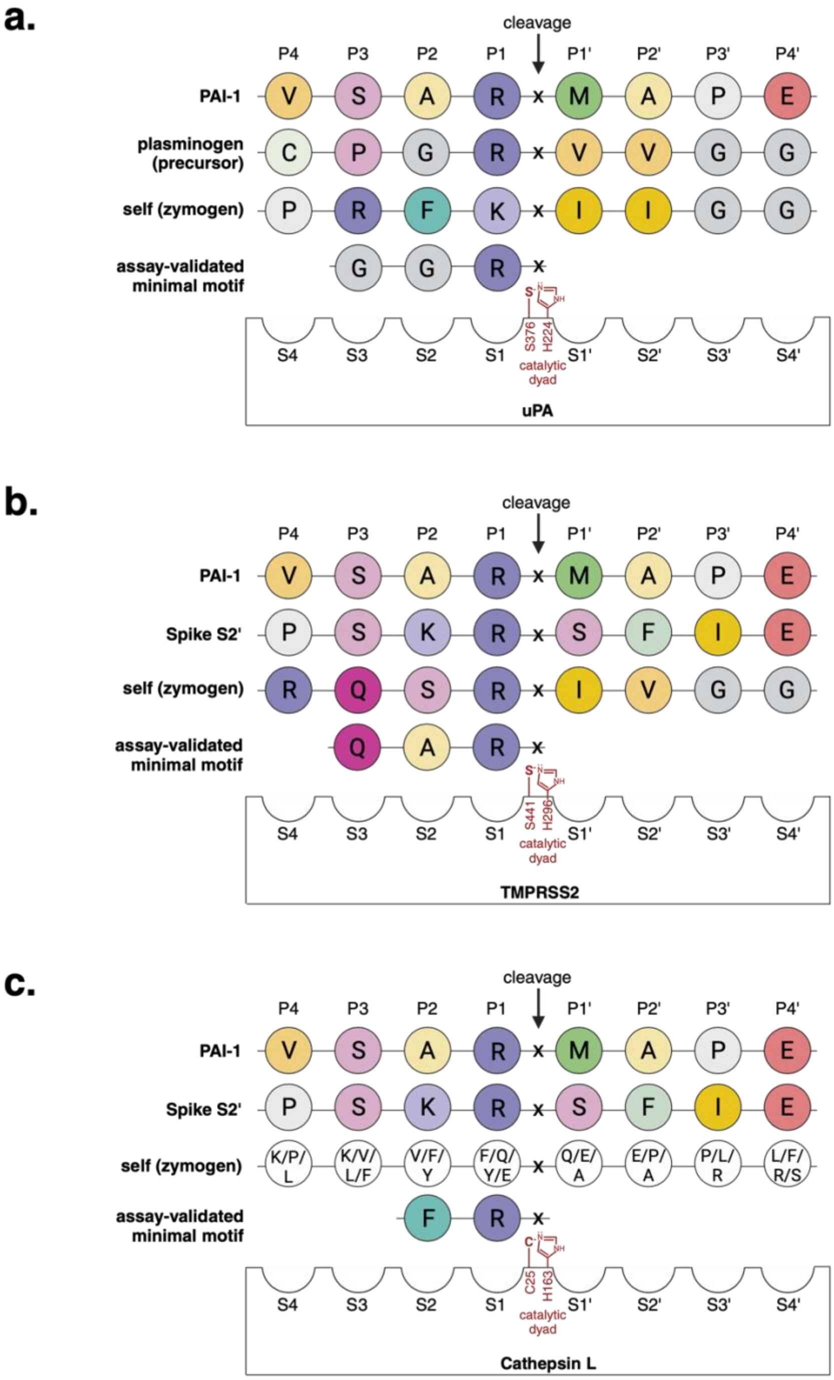


**Supplemental Figure 4. Schematic representation of substrate P4-P4’ motifs opposite proteases’ catalytic pockets S4-S4’ for uPA, TMPRSS2 and Cathepsin L.** Amino acids are colored according to their side chain chemistry (Unipro UGENE): basic (R, K) litmus blue with R being more basic and darker; acidic (E, D) litmus red with more acidic being darker; hydrophobic (I, L, V, A), yellow with intensity corresponding to hydrophobic character; sulfur-containing (C, M) green; aromatic (F, Y, W) in teal; polar (N, Q, S, T) magenta/pink with darker coloring for more polarity; non-polar glycine (G) in dark grey and proline (P) in light gray. Assembled from references[49-52] and UNIPROT.

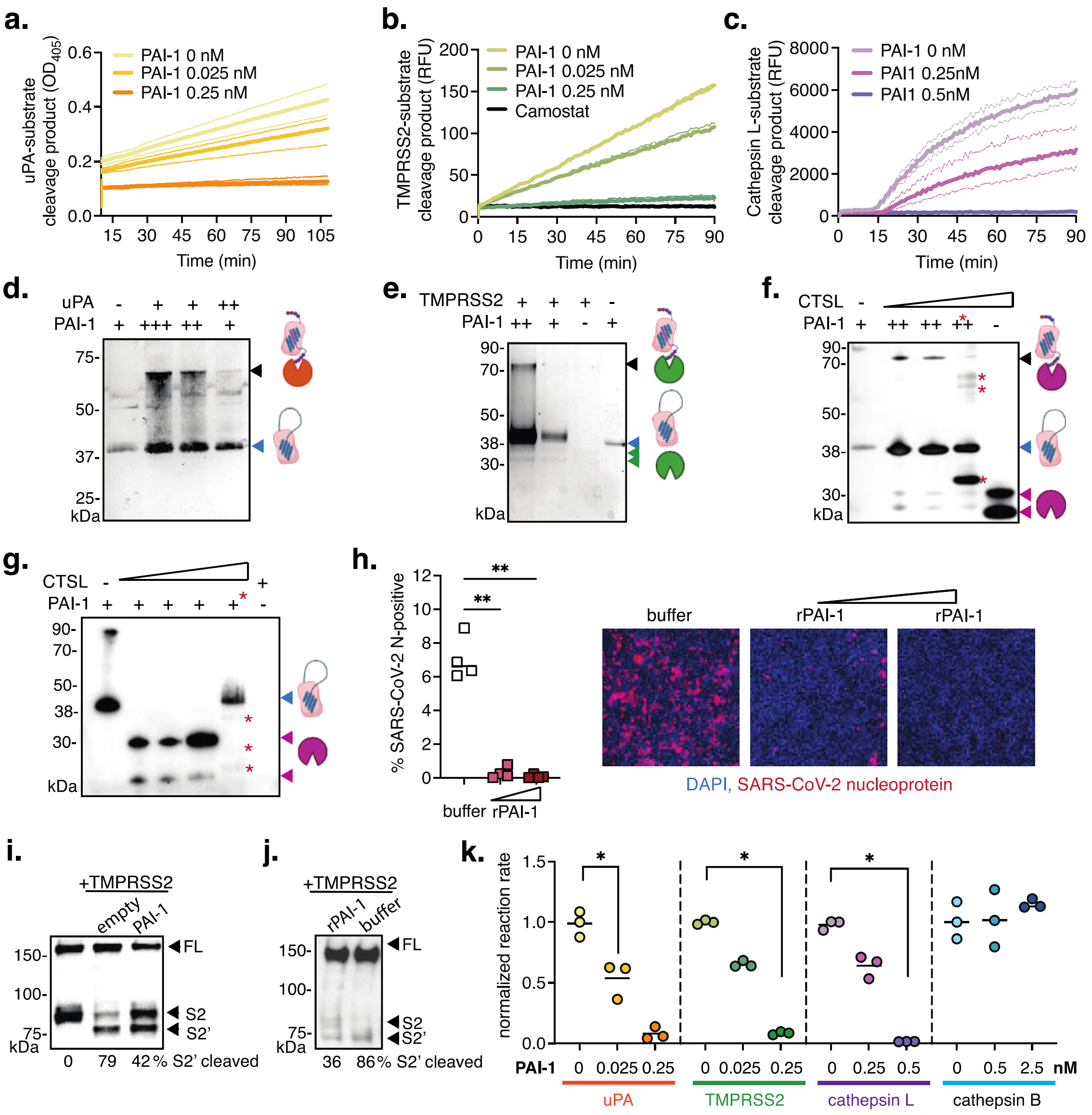


**Figure 4: PAI-1 impact on TMPRSS2 and cathepsins and its role in SARS-CoV-2 multi-cycle replication. a.** uPA protease activity assay with recombinant PAI-1 (Mean ± SD, n=3 replicates). **b.** TMPRSS2 protease activity assay with recombinant PAI-1 or camostat (Mean ± SD, n=3 replicates). **c.** Cathepsin L protease activity assay with recombinant PAI-1 at pH 6.5 (Mean ± SD, n=3 replicates). **d.** SDS-PAGE and silver stain of mixed recombinant active uPA (32 kDa, not visible) and PAI-1 (43 kDa). **e.** SDS-PAGE and silver stain of mixed recombinant active TMPRSS2 (31 kDa) and PAI-1 (43 kDa). **f.,g.** SDS-PAGE and silver stain of mixed recombinant cathepsin L (32 kDa) and PAI-1 (43 kDa) at pH 6.5 (f) or pH 5.5 (g). *Denotes use of PAI-1 inhibitor triplaxinin and PAI-1 cleavage products. **h.** SARS-CoV-2 multi-cycle infection in Calu-3 cells with extracellular addition of recombinant PAI-1 (48 hpi) by high-content microscopy. Statistical analysis by One-way ANOVA and Holm-Sidak's multiple comparison test, ** p<0.005. Representative images, DAPI (blue, nuclei), SARS-CoV-2 nucleoprotein (red). **i.** BHK cells co-transfected to express SARS-CoV-2 spike, TMPRSS2, and PAI-1. Spike S2 and 2' band intensities by western blot and densitometry. FL, full-length. **j.** BHK cells co-transfected to express SARS-CoV-2 spike and TMPRSS2, and rPAI-1 or buffer control added to the cell supernatant. Spike S2 and 2' band intensities by western blot and densitometry. FL, full-length. **k.** Normalized reaction rates for uPA, TMPRSS2, cathepsin L, and cathepsin B in the absence or presence of PAI-1 from *in-vitro* fluorometric assays. Statistical analysis by One-way ANOVA and Kruskal-Wallis test, * p<0.05.

88 pymol.bib

89 R Foundation for Statistical Computing, Vienna, Austria, 2014, https://www.r-project.org/foundation/

**Supplemental Table 1**

| SERPIN | F primer | R primer |
|---|---|---|
| A1 | TCTTCTTCTCCCCAGTGAGCATC | CAGGATTTCATCGTGAGTGTCAGC |
| A4 | GAGGCTAAACTCTTCCACACCAAC | TGACCAAATCCACAATCTTCCC |
| A5 | CAGACACTTTCCCCACCAACTTTAG | CCCTTCGTTTGCTTTGCCAC |
| A6 | CGGAGATGTGCTGGAGGAAATG | TGGACCACCTTTGATGACTTCAG |
| A7 | CCAACCAAATGCCACTCTCTACAAG | TGTTCTTATCTGGGGTCTCCACAG |
| A9 | TGTGCTCCAATCTACTGTGTGTCC | GCAAAGTCGGTGTTGAGGGAATAC |
| A10 | CTCACCCTTTGCTGACCTTAG | GCCCCTTTCATCAACTTCAATCAC |
| A11 | CCACAGAATCACACCCACCATTAC | GGCGAGAAGAAGATGTTTCCG |
| A12 | CCATTTTTCTGGCTGTTCTCCTC | CATCCTTGGACCTCGCTCAAAG |
| B1 | ATGGAGCAGCTGAGCTCAGC | CTAAGGGGAAGAAAATCTCCCC |
| B2 | AAATCCATTCATCCTTCCGCTC | CGCAGACTTCTCACCAAACAGC |
| B3 | CAGCATTAGGGATGGTCCTCTTAG | TCCTGTGGTGTTCTCTGTGACTTG |
| B4 | GCAAATGCTCCAGAAGAAAGTCG | GCCAATAGTCCCATCAGGAAATAGG |
| B5 | GGCAATGTCCTCTTCTCTCCAATC | TGTCACCTTTAGCACCCACTTGAG |
| B6 | CGCCGTAGAGAAGTCCAGAAAAC | CCAGAACCAGCCTTGTCAATGG |
| B8 | TGGAGCCAAGATTCTGTGC | TTTTGTGGTGCCTGATGAAG |
| B9 | GAAACTCACAGCCTGGACCAAG | AAATGCCGAAGCACAGATTCC |
| B10 | TCAGAAAGGACATCAACTCTTGGG | AGGGCGTTCACCAGAATCATCC |
| B11 | ATTCATTCCGAGTTTGGTGTCG | TTTGTCCCGTAGAGCCTGTTGG |
| B13 | TTTACTGTCACATCCGCCCCAG | GAAGAAAATCTGCCGAAGAAGAGG |
| C1 | GGAGAAGAAGGCAACTGAGGATG | AGAAAGTGGTAGCAAAGCGGG |
| D1 | CATCCTCAACGCCAAGTTCG | TGCCAACGGGTGCTATGAAG |
| E1 | GTGTTTCAGCAGGTGGCGC | CCGGAACAGCCTGAAGAAGTG |
| E2 | AAACGCACTTTCGTGGCAGC | ATCATTGGGGGCACTTGTCG |
| F1 | CAGCATTCTCCTTCTCGGTGTG | CACGGTCCTCTCTTCATCCAAGTAG |
| F2 | AACCAATGGGTGAAGGAGGC | AAGTGGATGGCGTTGAGGAGAAGC |
| G1 | GAGCCTCCTCAAATCCAAATGC | TTGTTGCGACCTTCCCTTCG |
| I2 | AAGTGTATGTTTCCCAAGTGACGC | GGATGTGTATGCCAGTTGATGTTG |
